# Reversible blood-brain barrier opening utilizing the membrane active peptide melittin *in vitro* and *in vivo*

**DOI:** 10.1101/2021.02.09.430012

**Authors:** Raleigh M. Linville, Alexander Komin, Xiaoyan Lan, Jackson G. DeStefano, Chengyan Chu, Guanshu Liu, Piotr Walczak, Kalina Hristova, Peter C. Searson

## Abstract

The blood-brain barrier (BBB) tightly controls entry of molecules and cells into the brain, restricting the delivery of therapeutics. Blood-brain barrier opening (BBBO) utilizes reversible disruption of cell-cell junctions between brain microvascular endothelial cells to enable transient entry into the brain. Development of BBBO techniques has been hindered by a lack of physiological models for *in vitro* study. Here, we utilize an *in vitro* tissue-engineered microvessel model to demonstrate that melittin, a membrane active peptide present in bee venom, supports BBBO. From endothelial and neuronal viability studies, we identify the accessible concentration range for BBBO. We then use a tissue-engineered model of the human BBB to optimize dosing and elucidate the mechanism of opening. Melittin and other membrane active variants transiently increase paracellular permeability via disruption of cell-cell junctions. In mice, we demonstrate a minimum clinically effective intra-arterial dose of 3 μM·min melittin, which is reversible within one day and neurologically safe. Melittin-induced BBBO represents a novel platform for delivery of therapeutics into the brain.

## 1. Introduction

The blood-brain barrier (BBB) remains a major roadblock for delivery of therapeutics to the brain [1, 2]. The ability to circumvent the BBB has the potential to enable the development of therapies for currently untreatable brain diseases [2]. Due to enriched expression of tight junction proteins by brain microvascular endothelial cells (BMECs), drug therapies have been largely limited to small lipophilic molecules that can passively diffuse across endothelial cell membranes [3]. While efflux transporter inhibition and hijacking of receptor-mediated transport systems have been extensively explored [4–6], these approaches have had limited clinical success. In contrast, blood-brain barrier opening (BBBO) involves reversible disruption of tight junctions to enable transient paracellular transport between endothelial cells. This approach has the advantage of allowing entry of any therapeutic or diagnostic agent into the brain over a limited period of time, followed by recovery of barrier function.

BBBO approaches can broadly be categorized as mechanical or chemical. Focused ultrasound (FUS) is a well-known method for mechanically-induced BBBO, involving disruption of tight junctions by distortion of microbubbles injected into systemic circulation [7, 8]. A well-known chemical method for BBBO is the intra-arterial (IA) injection of the hyperosmotic agent mannitol, which has been used in both preclinical models and clinical studies for several decades [9]. IA injection of mannitol results in endothelial cell shrinkage which can induce local transient disruption of tight junctions [9, 10]. This approach has been used to improve delivery of chemotherapeutics, stem cells, and viral vectors into the brain [11–15], however, widespread clinical use has been limited due to the lack of reproducibility [16–18]. Additionally, vasoactive compounds including bradykinin, bradykinin analogs, adenosines, arachidonic acid (AA), leukotrienes, and histamine have been explored for transient BBB opening, but have also had limited clinical success [19, 20].

Apart from hyperosmotic and vasoactive agents, little is known about other classes of molecules capable of intra-arterial BBBO. Previous studies have shown that systemic injection of membrane active peptides (MAPs) has the potential for permeabilizing biological barriers, including the BBB [21–23]. Importantly, MAP sequences can be engineered with customizable pharmacological properties (i.e. charge, hydrophobicity, amphipathicity) [24], presenting a large design space for optimizing clinical utility. To assess the potential for BBB opening, we selected melittin (MW 2847.5 Da), a small 26-amino acid MAP that is the main component (~50 wt.%) of honeybee venom [25] and several synthetic analogs [26]. Melittin and its analogs can disrupt plasma membrane integrity resulting in increased permeability of biological barriers [21, 22, 26–28]. However, systemic injection is extremely inefficient for drug delivery to the brain and can be associated with off-target effects in other organs. Therefore, we sought to determine concentrations and exposure times in which melittin, and its analogs, could support transient BBBO by modeling IA administration.

To answer this question, we took the unconventional approach of utilizing a tissue-engineered microvessel model of the human BBB to motivate *in vivo* experiments and elucidate mechanisms. 150 μm diameter microvessels were constructed using induced pluripotent stem cell (iPSC)-derived BMEC-like cells (dhBMECs), which recapitulate key features of the human BBB, including physiological permeability, cylindrical geometry, cell-matrix interactions, and shear stress [29, 30]. Melittin and its analogs were directly perfused into the microvessels, mimicking intra-arterial administration *in vivo*, and thus avoiding pharmacokinetic complexities associated with systemic delivery, which have been the focus of other human-on-a-chip *in vitro* systems [31]. This platform provides simultaneous real-time resolution on the dynamics of transport (e.g. leakage of different molecular weight solutes) and endothelial cell behavior (contraction, proliferation, apoptosis) following MAP treatment, which is challenging using conventional 2D systems.

Combining imaging of *in vitro* tissue-engineered microvessels and *in vivo* studies in mice, we describe BBBO using melittin, along with other synthetic analogs (**Fig. 1**). We determine peptide concentrations and exposure times that maintain viability of brain microvascular endothelial cells and neurons using a high-throughput microscopy-based cytotoxicity assay. We then determine dosing conditions for BBBO and elucidate the mechanism of disruption within our tissue-engineered microvessel model. Finally, we assess the BBBO following IA injection of melittin in mice using T1- and T2-weighted gadolinium-enhanced MRI, Evans blue leakage, and histology. We establish a minimum effective dose (MED) of melittin in mice which results in reversible BBBO within 24 hours, is neurologically safe, and 7-fold lower than doses required to open the human *in vitro* BBB. As animal models are poor predictors of clinical translation, these results highlight potential species-to-species differences that will need to be further explored prior to clinical application. Overall, our work highlights tissue-engineered models as a promising tool for translation of drug delivery strategies to clinical application, and specifically develops MAPs as a novel class of compounds for transient BBBO.

**Fig. 1.**
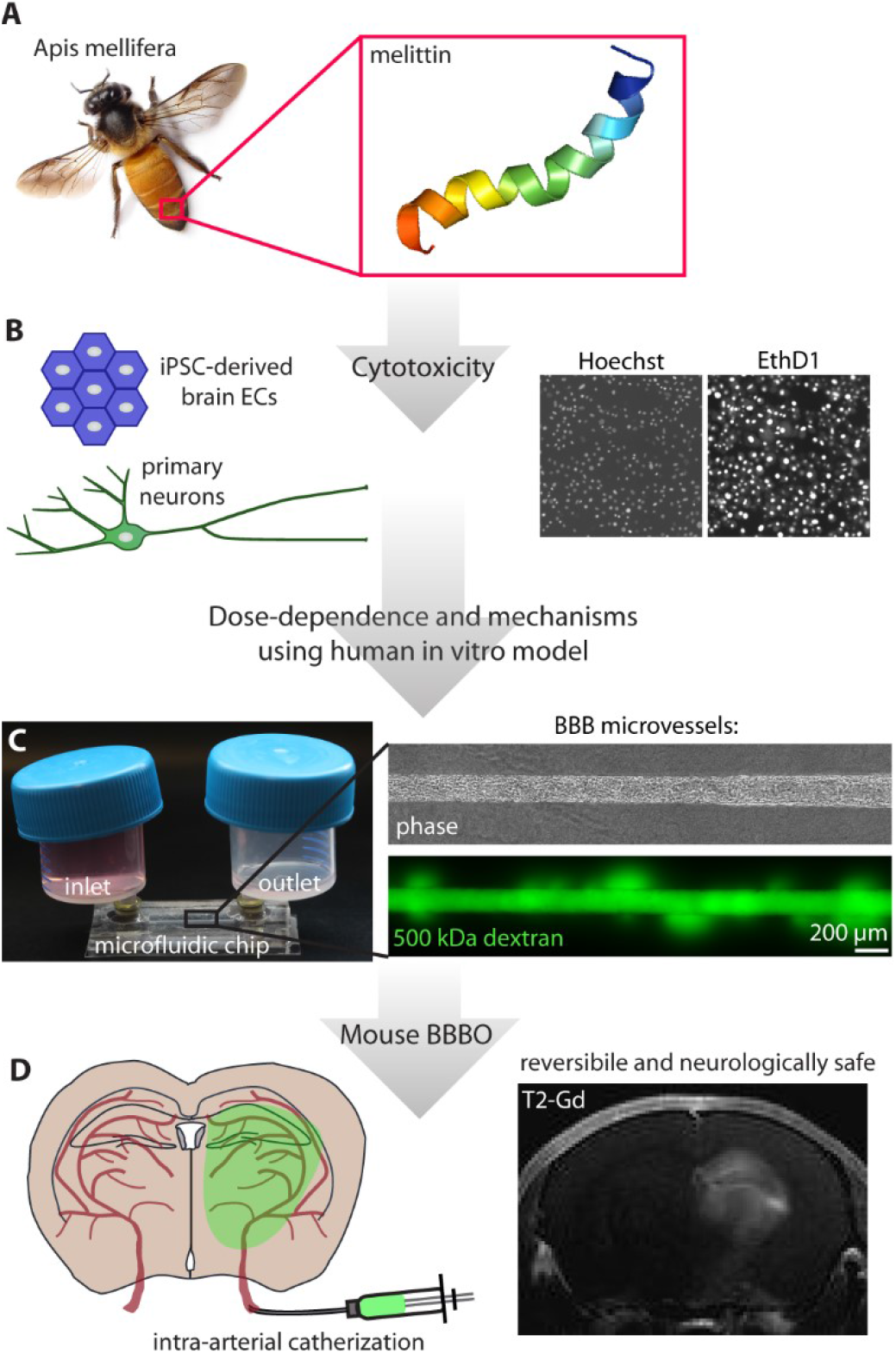
Development of membrane-active peptide (MAP)-induced intra-arterial blood-brain barrier opening. **(A)** Melittin is a 26 amino acid MAP and the main component of honeybee (*Apis mellifera*) venom. The inset shows a 3D model of melittin structure from the RSCB Protein Data Bank (doi: 10.2210/pdb6dst/pdb). **(B)** Cytotoxicity assays used to determine doses of melittin that minimally compromise brain microvascular endothelial cell and neuronal viability. **(C)** Tissue engineered human microvessel model used to determine dose dependence and mechanisms of BBBO. **(D)** Application of MAP as an intra-arterial injection into the brain supports reversible and neurologically safe BBBO.

## 2. Methods

### 2.1. Cell culture

Induced pluripotent stem cell (iPSC)-derived BMEC-like cells (dhBMECs) mimic the barrier function of the human BBB [32–34]. dhBMECs were differentiated as previously reported [29, 35, 36]. The BC1 iPSC line derived from a healthy individual was used for all experiments [37]. After eleven days of differentiation, cells were detached using Accutase (A1110501, Thermo Fisher Scientific), and seeded onto transwells, glass slides, or type I collagen microchannels coated overnight with 50 μg mL^−1^ human placental collagen IV (Cat. no. C5533, Sigma-Aldrich) and 25 μg mL^−1^ fibronectin from human plasma (Cat. no. F1141, Sigma-Aldrich). Cells have previously been characterized for expression of tight junction proteins, efflux pumps, and nutrient transporters as well as measurements of barrier function including transendothelial electrical resistance (TEER) and solute permeability [32, 35]. Here, cells with TEER of at least 1500 Ω cm^2^ were used for all experiments. After detachment, cells were cultured in BBB induction medium: human endothelial cell serum-free medium (HESFM) (Cat. no. 11111044, Thermo Fisher Scientific) supplemented with 1% human platelet poor derived serum (Cat. no. P2918, Sigma-Aldrich), 1% penicillin-streptomycin (Cat. no. 15140122, Thermo Fisher Scientific), 2 ng mL^−1^ bFGF (Cat. no. 233FB025CF, Fisher Scientific), and 10 μM all-trans retinoic acid (Cat. no R2625, Sigma-Aldrich). After 24 hours, cells were further cultured in BBB maintenance medium: HESFM supplemented with 1% human platelet poor derived serum and 1% penicillin-streptomycin.

Human cortical neurons (HCNs, CRL-10742) were obtained from ATCC (Manassas, VA, USA). HCNs were cultured in DMEM (Cat. no. 31600091, Thermo Fisher Scientific), adjusted to contain 1.5 g L^−1^ sodium bicarbonate, 4.5 g L^−1^ D-glucose, and supplemented with 10% heat-inactivated FBS (Cat. no. F4135, Sigma-Aldrich), following ATCC recommendations.

### 2.2. Peptide preparation, characterization, and exposure

Melittin (melittin-COO^−^), melittin-CONH_2_, scrambled melittin-CONH_2_, MelP3, and MelP5 were synthesized at > 95% purity by Bio-Synthesis Inc (Lewisville, TX, USA). Peptide sequences and chemical structures are shown in **Table S1** and **Fig. S7**. Melittin with a free carboxy C-terminus (melittin-COO^−^) was used for all experiments to maximize water solubility and hence accessible concentration range, unless specified. Peptide stocks were prepared by resuspending lyophilized peptide powders in MilliQ H_2_O, following aliquoting and storage at −20 °C. Peptide stock aliquots were used for a maximum of three freeze-thaw cycles. Peptide stock solutions were diluted in BBB maintenance medium and prepared fresh prior to experiments.

Circular dichroism spectra of 5 μM melittin and scrambled melittin-CONH_2_ were measured in 20 mM sodium phosphate buffer with 20 mM SDS (pH 7.4) using Jasco J-710 spectropolarimeter (Easton, MD, USA). The spectra were measured at a scanning speed of 100 nm min^−1^, data pitch 0.1 nm, response 4 s, and bandwidth 1 nm from 280 nm to 180 nm.

### 2.3. Transwell assay

Corning transwell supports (Cat. no. 3470) were coated with 25 μg mL^−1^ fibronectin and 50 μg mL^−1^ collagen IV overnight at 37 °C. DhBMECs were seeded on transwells in BBB induction medium at 10^6^ cells cm^−2^. On day 1 after seeding, the medium was changed to the BBB maintenance medium after one wash with the maintenance medium. On day 2, TEER measurements were performed using an EVOM-2 and STX-100 electrodes (World Precision Instruments, Sarasota, FL). Monolayers with TEER > 1500 Ω cm^2^ were used for the permeability experiments. All TEER measurements are reported after subtracting the value for a blank transwell correcting for the membrane area. For transwell permeability measurements, cell monolayers were washed with BBB maintenance medium, followed by the addition of 500 kDa dextran-fluorescein (1.2 μM) to the apical side of the transwell with or without 5 μM melittin. The transwells were incubated, while rocking, for 90 min at 37 °C, 5% CO_2_. After the permeability experiments, the TEER across monolayers was measured. Dextran fluorescence in the basolateral and apical chambers was measured using the Synergy H4 plate reader (Biotek Instruments Inc., Winooski, VT) and was converted to a dextran concentration using dextran-fluorescein standard curves. Dextran concentrations in the basolateral chambers were less than 10% of the input concentration (average = 0.1%), confirming that transport was in the linear regime [38]. The apparent permeability (P_app_) was calculated from:

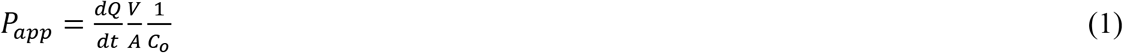

where *dQ*/*dt* is the increase in dextran concentration in the basolateral chamber with time, *V* is the volume of the basolateral chamber, *A* is the surface area of the transwell support, and *C*_o_ is the initial dextran concentration in the apical chamber. The permeability experiments were performed with dhBMECs from three independent iPSC differentiations.

### 2.4. Immunocytochemistry

Antibodies used in the study are summarized in **Table S2**. For staining of confluent monolayers of dhBMECs, LabTek 8-chamber slides (Cat. no. 155411, Thermo Fisher Scientific) were coated with 25 μg mL^−1^ fibronectin and 50 μg mL^−1^ collagen IV overnight at 37 °C. dhBMECs were then seeded onto the slides in BBB induction medium at 5×10^5^ cells cm^−2^. On day 1 after seeding, dhBMECs were washed twice with BBB maintenance medium and incubated in the medium overnight. On day 2, the cells were washed twice with BBB maintenance medium and treated with 1.5 μM, 5 μM, and 10 μM of melittin, diluted in the medium, for 2 min, 5 min, 10 min, or 90 min. The resulting melittin dosages, calculated as the product of melittin concentration and incubation time, were 3 μM·min, 25 μM · min, 100 μM · min, and 450 μM·min. A negative control without melittin was included with every experiment. After the melittin incubation, the cells were washed twice with 1xPBS azide and fixed in 3.7% paraformaldehyde solution for 10 min or methanol for 15 minutes. The fixed cells were washed with 1xPBS azide and incubated in blocking solution (10% goat serum, 0.3% TritonX-100 in 1xPBS azide) at 4°C for 1-2 days. The cells were incubated with primary antibodies for 2 hours at room temperature. After the incubation, the cells were washed three times 5-minute each with PBS azide, and incubated with secondary antibodies at 1:200 dilution for one hour at room temperature, followed by three 5-minute washes. Afterwards, the cells were incubated with Fluoromount G, containing DAPI (Cat. no. 00-4959-52, Thermo Fisher Scientific). Epifluorescence images were taken with an inverted microscope Nikon Eclipse TiE using 40x water immersion objective and NIS Advanced Research software (Nikon, Minato, Tokyo, Japan).

### 2.5. Cytotoxicity assay

DhBMECs were seeded on polystyrene 96-well plates, coated with 25 μg mL^−1^ fibronectin and 50 μg mL^−1^ collagen IV, at 2×10^5^ cells cm^−2^ in BBB induction medium. On day 1, the cells were washed twice with BBB maintenance medium with gentle pipetting up-and-down to remove non-adherent cells and incubated in the medium overnight. HCN2 cells were seeded in the tissue culture-treated 96-well plate at 3×10^3^ cells cm^−2^ in HCN2 cell culture medium and cultured for two days at 37 °C, 5% CO_2_, with the medium change on day 1 after seeding. The cytotoxicity assay was performed on day 2 after seeding. Briefly, cells were incubated with two dyes: a membrane-impermeable Ethidium homodimer-1 (EthD1, L3224, Thermo Fisher Scientific) to assess cell viability, and Hoechst 33342 nuclear stain (Cat. no. 62249, Thermo Fisher Scientific), and imaged on an epifluorescence microscope. The cells were washed with BBB maintenance medium followed by incubation with 200 μL of 2 μg mL^−1^ Hoechst 33342 and 10 μM EthD1 solution for 15 - 30 min at 37 °C, 5% CO_2_. After the incubation, 100 μL of the solution was replaced with Hoechst 33342 and EthD1 solutions containing melittin using a multichannel pipettor, resulting in final concentrations of 0.75 μM, 5 μM, and 10 μM melittin. After the addition of treatment solutions, large 6 x 6 epifluorescence images were collected every ~3.5 min per condition using a 20x objective and a Nikon Eclipse TiE microscope, equipped with 37 °C chamber, for 60 – 90 min. The out-of-focus portions of images were manually removed using ImageJ. Afterwards, the images were analyzed using a custom pipeline in CellProfiler 3.1.8 to count Hoechst-positive objects (all cells) and EhtD1-positive objects (dead cells) [39]. The fraction of viable cells (*F*) was calculated from:

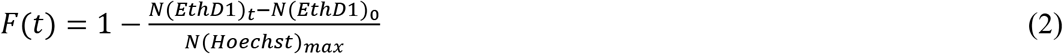

where *N(EthD1)*_*t*_ is the number of EthD1-positive objects at time t, *N(EthD1)*_*0*_ is the number of EthD1-positive objects at time t = 2 min (initial time of imaging), *N(Hoechst)*_*max*_ is the maximum number of Hoechst-positive objects counted in an image over the imaging time-course. For dhBMECs, the experiments were repeated a total of five times with two independent iPSC differentiations. In these experiments, to avoid monolayer disruption media removal steps were performed by gentle pipetting and not by aspiration. For HCN2 cells, the experiments were performed a total of three times on day 2 and day 3 after seeding.

### 2.6. Fabrication of tissue-engineered microvessels

Three-dimensional blood-brain barrier microvessels were fabricated as previously reported (**Fig. S3A,B**) [29]. 150 μm diameter and 1 cm long channels were patterned in 7 mg mL^−1^ type I collagen (Corning) using nitinol wire (Malin Co.) housed within polydimethylsiloxane (PDMS; Dow Corning). Channels were cross-linked by incubation in 20 mM genipin (Wako Biosciences) for 2 hours to increase cell adhesion [36]. dhBMECs were suspended in BBB induction medium at a concentration of 5 × 10^6^ cells mL^−1^ and seeded into channels. Microvessels were perfused using BBB induction medium for 24 hours and then switched to BBB maintenance medium. During cell seeding and the first day of perfusion, 10 μM ROCK inhibitor Y27632 (ATCC) was supplemented to increase adhesion [36]. An average shear stress of ~ 2 dyn cm^−2^ was maintained with a reservoir height difference (Δh) of 3 cm, corresponding to a volumetric flow rate of ~250 μL h^−1^ (**Fig. S3D**). This shear stress was estimated from the Poiseuille equation: *τ* = *μQ* /2π*d*^3^ where *μ* is the dynamic viscosity, *Q* is the volumetric flow rate, and *d* is the microvessel diameter.

### 2.7. Live-cell imaging of tissue-engineered microvessels

Live-cell imaging was conducted to study dynamic changes in permeability and microvessel structure. Microvessels were imaged using a 10x objective on an inverted microscope (Nikon Eclipse TiE) maintained at 37 °C and 5% CO_2_ (**Fig. S3C**). Epifluorescence illumination was provided by an X-Cite 120LEDBoost (Excelitas Technologies). Phase contrast and fluorescence images were acquired every two minutes with the focal plane set to the microvessel midplane. The entire image consisted of ten adjacent imaging frames. Microvessels were imaged for 20 minutes before perfusion with fluorescent solutes, and 90 minutes after addition of fluorescent solutes and melittin. A dextran solute panel comprised of 200 μM Cascade Blue-conjugated 3 kDa dextran (Cat. no. D7132, Thermo Fisher Scientific), 16 μM Alexa Fluor647-conjugated 10 kDa dextran (Cat. no. D22914, Thermo Fisher Scientific), and 1.2 μM fluorescein-conjugated 500 kDa dextran (Cat. no. D7136, Thermo Fisher Scientific) was prepared in BBB maintenance medium. Fluorophores were independently excited and the respective emissions were collected using: Nikon UV-2E/C for 3 kDa dextran (50 ms exposure), Chroma 41008 for 10 kDa dextran (20 ms exposure), and Nikon B-2E/C for 500 kDa dextran (50 ms exposure).

### 2.8. Analysis of BBB opening

Image processing was performed using ImageJ (NIH) to quantify BBB opening (**Fig. S3**). First, images were cropped to a standard size of 6 mm x 0.6 mm. To determine the spatial heterogeneity of BBB permeability, microvessels were sectioned into 10 regions of interest (ROIs), each a 0.6 mm x 0.6 mm square (**Fig. S3A-i**). For each ROI, the integrated pixel density was plotted over time (**Fig. S3A-ii**). The permeability (cm s^−1^) was calculated using *P* = (*r*/2)(1/*ΔI*)(*dI*/*dt*), where *r* is the vessel radius, *ΔI* is the increase in fluorescence intensity upon perfusion of the fluorophore, and (*dI*/*dt*) is the rate of increase in fluorescence intensity as solute exits into the gel [40]. The average permeability (*P*_avg_) was calculated from a linear least squares fit to *dI*/*dt* over 90 minutes, while the instantaneous permeability (*P*_inst_) was determined from a linear least squares fit to *dI*/*dt* in 10-minute increments upon melittin perfusion (**Fig. S3A-iii**). To visually display permeability dynamics, heatmaps were generated from 90 instantaneous permeability values across the 10 ROIs (x-axis) and nine 10-minute periods (y-axis).

To study the dynamics of solute leakage induced by melittin, fluorescence images were converted to binary images using the Auto Threshold function (plugin v1.17; Huang2 method) [41] in ImageJ (**Fig. S3B**). Under baseline conditions, thresholding results in the lumen designated as white (1) and extracellular matrix (ECM) as black (0). Focal leaks due to melittin exposure appear as white plumes that extend from the lumen into the ECM that emerge (or disappear) over time. From the images, individual focal leaks were manually counted over time. Leaks emerging from the sides of the image frame were not counted. To quantify the duration of a BBB opening event, we identified the time at which the last focal leak appeared, termed the focal leak reversibility time constant (τ_focal leaks_).

### 2.9. Analysis of cell behavior

To monitor cell dynamics, cell loss and proliferation events were counted manually. Rates of cell loss (% h^−1^), division (% h^−1^), turnover (% h^−1^), and cell loss at the site of a focal leak (%) were quantified from phase contrast imaging sequences, as reported previously [29, 42]. Briefly, the total number of endothelial cells were counted manually for each microvessel plane, utilizing a region approximately 100 μm wide corresponding to about a fifth of the channel circumference. Cell loss was identified by observing the removal of a cell from the microvessel. Cell loss from the monolayer occurred through two processes: (1) apoptotic loss, in which a cell constricts, releases its contents into the vessel lumen, and is then removed from the monolayer over a period of 20 – 60 minutes, and (2) non-apoptotic loss, in which a cell swells and disappears from the monolayer over a period of 2 – 6 minutes. Cell division was identified by observing cell compression, alignment of chromosomes, and the formation of daughter cells. The rate of turnover (% h^−1^) is the difference between the rate of cell division and the rate of cell loss. A cell loss event was identified to be at the site of a focal leak if it occurred within < 50 μm and < 4 minutes of the occurrence of a focal leak. For a limited set of experiments, the WTC iPSC line [43] with GFP-tagged zona occludens-1 (Allen Cell Institute) was used to facilitate live-cell imaging of tight junction organization.

### 2.10. Confocal imaging of tissue-engineered microvessels

Confocal z-stacks (0.4 μm in thickness) at 40x magnification were obtained on a swept field confocal microscope system (Prairie Technologies). Illumination was provided by an MLC 400 monolithic laser combiner (Keysight Technologies). Complete reconstruction of the microvessel lumen required approximately four hundred slices. Confocal imaging was conducted before, during, and after exposure to 50 μM·min melittin treatment.

### 2.11. Intraarterial injection of melittin in mice

Animal procedures were approved by The Johns Hopkins Animal Care and Use Committee and were performed in accordance with ARRIVE guidelines. The surgical procedures for carotid artery catheterization were performed as described previously [44]. Briefly, male C57BL/6J mice (*n* = 43, 6 – 8 weeks old, 20 – 25 g, Jackson Laboratory) were anesthetized with 2% isoflurane. The common carotid artery (CCA) bifurcation was exposed and the occipital artery branching off from the external carotid artery (ECA) was coagulated. The ECA and the pterygopalatine artery (PPA) were temporarily ligated with 4–0 silk sutures. A temporary ligature using a 4–0 suture was placed on the carotid bifurcation and the proximal CCA was permanently ligated. A microcatheter (PE-8-100, SAI Infusion Technologies) was flushed with 2% heparin (1,000 U mL^−1^, heparin sodium, Upjohn), inserted into the CCA via a small arteriotomy, and advanced into the internal carotid artery (ICA).

To explore the optimal concentration of melittin for blood-brain barrier opening (BBBO), 1 μM, 3 μM and 5 μM melittin prepared in saline was infused through the ICA catheter by a syringe pump (11 PLUS, Harvard Apparatus Inc.). For each animal, 150 μL of solution was infused at a rate of 150 μL min^−1^. To evaluate BBB status, 0.2 mL of 2% w/v Evans Blue in saline was subsequently administrated through tail vein injection immediately post-melittin infusion and in a separate cohort after 24 h and 7 days. The brains were harvested immediately after Evans Blue injection, sectioned in brain slicer matrix (Zivic Instruments) to produce 1 mm slabs. After identification of the lowest concentration of melittin that was effective for BBBO, an additional group of animals (*n* = 5) was observed for 30 days to comprehensively assess safety of the procedure. Neurological deficits were assessed by visual inspection to confirm that mice displayed no paralysis, seizure, or rotation.

### 2.12. MRI and histological assessment of BBBO

Animal anesthesia was maintained at 1.5 - 2% during the MRI scan. Mice were placed on a water-heated animal bed equipped with temperature and respiration control. Respiration was monitored and maintained at 30 – 60 min^−1^. All MRI experiments were performed on a Bruker 11.7T MRI scanner. Baseline T2 (TR/TE = 2,500/30 ms), susceptibility-weighted imaging (SWI) (TR/TE = 378/8 ms), T1 (TR/TE 350/6.7 ms)-weighted, dynamic gradient echo echo-planar imaging (GE-EPI, TR/TE 1250/9.7 ms, field of view (FOV) = 14 × 14 mm, matrix = 128 x 128, acquisition time = 60 s and 24 repetitions) and dynamic T1 imaging (TR/TE 2000/3.5ms, field of view = 14×14 mm, acquisition time = 8 s and 20 repetitions) images of the brain were acquired. The microcatheter was connected to a syringe mounted on an MRI compatible programmable syringe pump (PHD 2000, Harvard Apparatus Inc.). Gadolinium (Gd; Prohance) dissolved in saline at 1:50 was infused intra-arterially at the rate of 150 μL min^−1^ under dynamic GE-EPI MRI for visualization of perfusion territory. Next 1, 3 or 5 μM melittin mixed with Gd (50:1) was administered intra-arterially for one minute at an injection speed of 150 μL min^−1^. For dynamic monitoring of BBBO, T1 scans were acquired during and after a 1-minute infusion of melittin. For assessment of the BBBO status, T1-weighted image was collected after melittin infusion and repeated at 24 h and 7 days. To quantify BBB status, T1 intensities in the cortex, hippocampus, and thalamus/hypothalamus were compared between ipsilateral and contralateral hemispheres. T1 hyperintensity was corrected for possible asymmetry using pre-contrast T1 images and normalized as the ratio of ipsilateral to contralateral intensity. Additionally, the area of BBBO (mm^2^) was calculated by conversion of T1 images to binary images using the Auto Threshold function (plugin v1.17; Maximum Entropy method) [45] in ImageJ. T2-weighted and SWI images were acquired at the same time points for detection of brain damage and microhemorrhage. Subsequently, animals were sacrificed for further histological assessment.

To determine the area and region of the perfusion territory, we first created contrast enhancement maps at the time when contrast enhancement was maximum. As Gd causes negative contrast under T2 imaging, we calculated the contrast enhancement as (*SI*^pre^-*SI*^post^)/*SI*^pre^, where *SI* is the signal intensity of an MR image. Histograms were obtained for all pixels in the brain region in a contrast enhancement map and further fitted to two Gaussian distributions, assuming there were two dominant populations for non-contrast enhanced and contrast-enhanced pixels. The cutoff value to separate the two populations was then determined. The fraction of contrast enhanced areas was calculated by the ratio of the number of pixels, whose values are larger than the cut-off value, to the total number of pixels in the brain. A similar method was used to determine the BBBO territory. In brief, each post-contrast image was first fitted into two Gaussian distributions, and the cut-off of signal intensity was determined. Data analysis and the calculation of area fraction were performed using custom-written MATLAB scripts.

24 hours and 7 days after melittin treatment, animals were trans-cardially perfused with 5% sucrose in deionized water and then 4% paraformaldehyde in phosphate buffered saline. The brains were cryopreserved in 30% sucrose and cryo-sectioned in 30 μm slices. Primary antibodies used in the study are described in **Supplemental Table 2**. Goat anti-rabbit Alexa Fluor-488 or Alexa Fluor-594 (1:250, Life Technologies) were used as secondary antibodies for fluorescence assessment using an inverted microscope (Zeiss, Axio Observer Z1). For demyelination assessment, eriochrome cyanine R staining was performed as described previously [46], with assessment using bright-field microscopy. Quantitation of immunohistochemistry (based on relative fluorescence intensity) and MRI analysis (based on intensity) were performed using Image J. For each mouse, three ROIs of the ipsilateral and contralateral BBBO regions were drawn, respectively.

### 2.13. Statistical analysis

Melittin doses were randomly assigned in both microvessel and mouse experiments, and data analysis was performed without knowledge of the dose. Data were excluded if microvessel imaging planes were not in optical focus to adequately conduct analysis or if iPSC differentiations produced endothelial cells with transendothelial electrical resistance below 1500 Ω cm^2^. Animals were excluded if their death was related to the complications of surgery, including bleeding and hyperanesthesia.

All statistical analysis was performed using Prism ver. 8 (GraphPad). Metrics are presented as means ± standard error of the mean (SEM). The principle statistical tests used were a student’s unpaired t-test (two-tailed with unequal variance) for comparison of two unpaired groups, a student’s paired t-test (two-tailed with unequal variance) for comparison of two paired groups, and an analysis of variance (ANOVA) for comparison of three or more groups. For ANOVA tests, reported *p* values were multiplicity adjusted using a Tukey test. Linear regression of metrics across experimental parameters was conducted using least squares fitting with no constraints. An *F* test was used to determine if linear regression produced a statistically significant non-zero slope; *p* values were not calculatable in the event of perfect fits. An extra sum-of-squares *F* test was used to compare slopes between linear fits of dose-dependent behaviors. Differences were considered statistically significant for *p* < 0.05, with the following thresholds: * *p* < 0.05, ** *p* < 0.01, *** *p* < 0.001.

## 3. Results

### 3.1. Melittin causes chemically-induced BBB opening

Melittin induces disruption in confluent monolayers of stem-cell derived brain microvascular endothelial cells (dhBMECs) at 5 μM concentrations (**Fig. S1**). After 24 hours of exposure transendothelial electrical resistance is substantially decreased (*p* = 0.009), permeability is increased (*p* < 0.001) and holes become visible in the endothelium. In contrast, mannitol induces BBBO above a concentration of about 1.1 M, close to the solubility limit of around 1.4 M [9]. Therefore, even with IA injection, the concentration window for mannitol-induced BBBO is extremely limited and small decreases in concentration resulting from mixing with the blood due to collateral circulation can affect reproducibility and territories of BBBO [47, 48]. In contrast, the solubility of melittin is about 350 μM, more than 70-fold larger than the concentrations for causing irreversible damage (5 μM), providing a broad window for establishing reversible and non-cytotoxic conditions.

### 3.2. Cell viability in response to melittin dosing

Chemically-induced BBBO relies on exposing brain microvascular endothelial cells (BMECs) to molecules that induce cell stress. Therefore, reversible BBBO is a balance between inducing disruption of cell-cell junctions to enable paracellular transport, while minimizing cell death and maximizing the ability of the BMECs to recover and reform junctions after exposure. We hypothesized that the upper limit of the concentration range for reversible BBBO is dependent, in part, on the limit of viability of BMECs exposed to melittin. We first determined the viability of dhBMECs and human cortical neurons (HCNs) exposed to melittin. To accurately assess the limits of viability, we developed a sensitive imaging-based live-cell assay to measure the fraction of ethidium homodimer 1 (EthD1)-positive (i.e. dead) cells (**Figs. 2 and S2**). The assay involves time-course imaging of cells in medium containing melittin, nuclear stain (to count all cells), and EthD1 (to count dead cells) and using automated image analysis to count the total number of cells and dead cells over time. Upon exposure to 5 μM melittin, the viability of dhBMECs and HCNs was largely unaffected for about 20 minutes but then decreased sigmoidally (**Fig. 2A**). This suggests that there is a delay in the cytotoxic or membrane-permeabilizing effect of melittin on cells. The time at which the viability reached 50% (τ_1/2_) was ~30 minutes for HCNs compared to ~40 minutes for dhBMECs (*p* = 0.015) (**Fig. 2B**). Similarly, we observed shorter τ_1/2_ for HCNs when compared to dhBMECs, across other melittin concentrations (**Fig. S2)**. Although cortical neurons appear to be more sensitive to melittin exposure than dhBMECs, in the brain they are shielded from compounds in circulation by the BBB and thus are expected to be exposed to lower melittin concentrations than dhBMECs.

**Fig. 2.**
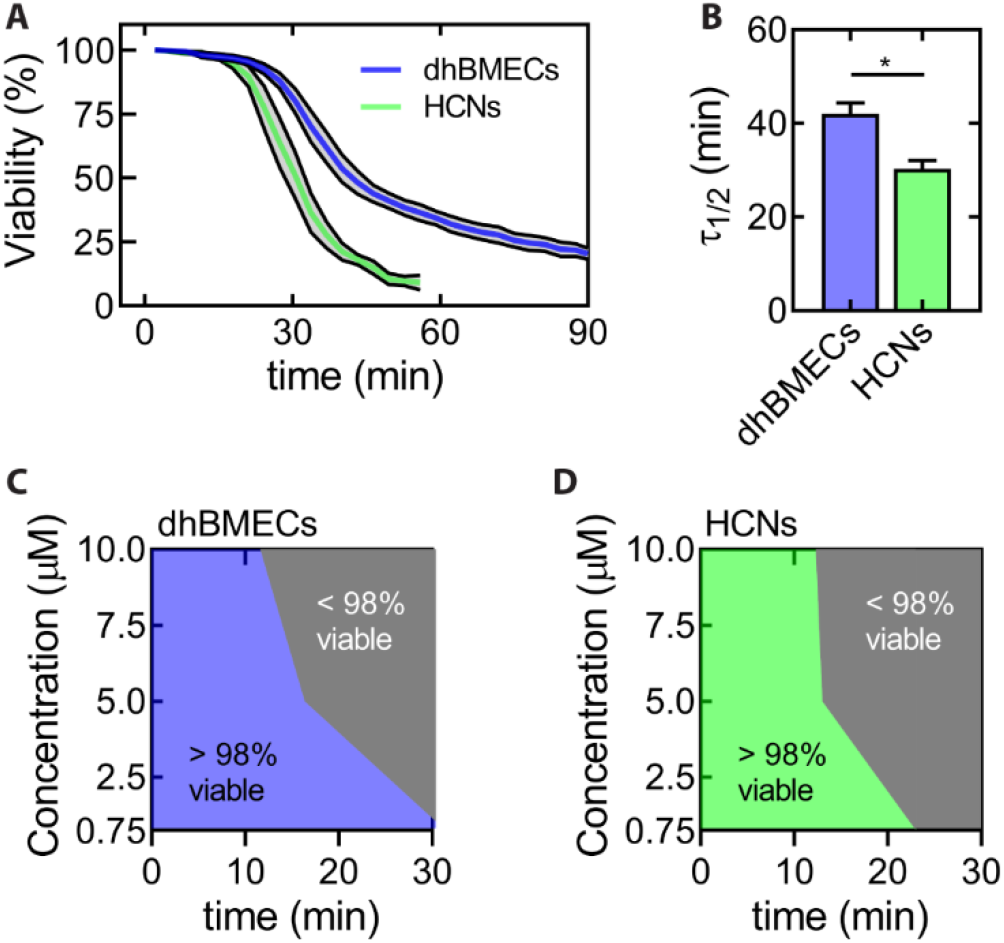
Melittin-induced cytotoxicity of derived human brain microvascular endothelial cells (dhBMECs) and human cortical neurons (HCNs). **(A)** Time course of cell viability for dhBMECs and HCNs incubated with 5 μM melittin. **(B)** Viability half-life for both cell types. Data represent mean ± SEM. **(C, D)** Phase diagrams of 98% cell viability for dhBMECs and HCNs, respectively. The phase diagrams show melittin concentrations and perfusion times which result in lower or greater than 98% cell viability. n = 5 for dhBMECs and n = 3 for HCNs. Statistical significance was calculated by a student’s unpaired t-test. * *p* < 0.05. Data presented as means ± SEM.

To identify the limit of cell viability on exposure to melittin (defined as the highest concentration where the viability is greater than 98%), we constructed phase diagrams (**Figs. 2C, D**). High cell viability can be maintained by short exposure (≤ 10 min) to 10 μM melittin or longer exposure (≥ 20 min) to 0.75 μM melittin. The relationship between concentration and exposure time that leads to 98% viability is not linear. Decreasing the concentration from 10 μM to 5 μM melittin provides only a slight extension in exposure time for maintaining low cytotoxicity. A short perfusion time is more clinically relevant for IA injection, and hence we proceeded to investigate the effect of melittin on BBBO for short perfusion times, as they result in low toxicity. Since a dose of 100 μM·min decreases dhBMEC and HCN viability by < 2%, we hypothesized that enhancements in permeability could be reversible below this dose.

### 3.3. BBBO in a tissue-engineered microvessel model

To assess the extent of BBBO and potential for recovery, we utilized a tissue-engineered BBB model [29, 30]. 150 μm diameter channels patterned in cross-linked collagen I were seeded with dhBMECs and maintained under a shear stress of 2 dyne cm^−2^ (**Fig. S3**). Microvessels were perfused with melittin and fluorescently-labeled dextran under live-cell imaging to monitor changes in permeability and cell behavior (**Fig. S4**). This model recapitulates the physiological permeability of the human BBB by restricting paracellular permeability of 500 kDa dextran under baseline conditions (**Supp. Video 1**). During melittin exposure, barrier function is transiently lost and leakage of 500 kDa dextran is observed (**Fig. S5 and Supp. Video 2**).

Having identified a melittin dose of 100 μM·min as the upper limit to maintain neuronal and endothelial viability, we perfused BBB microvessels with various melittin concentrations (1.5, 5 and 10 μM) for different exposure durations (2, 5 and 10 minutes), representing a wide range of doses from 3 to 100 μM·min (**Fig. 3**). To monitor changes in barrier function, dyes were perfused during and after melittin dosing for a total of 90 minutes (**Fig. 3A**). Low doses of melittin did not induce BBBO as evident from images recorded during perfusion with 500 kDa dextran, which resembled those of control BBB microvessels (**Figs. 3B-I and S5A**). Intermediate doses of melittin induced BBBO and resulted in recovery during the 90-minute imaging period (**Fig. 3B-ii**). High doses of melittin induced sustained BBBO and, although opening was not reversed during the 90 minute imaging period following administration (**Fig. 3B-iii**), barrier function was regained 24 hours later. To visualize the spatial and temporal heterogeneity of BBBO, heatmaps were generated for each melittin dose (**Figs. 3C and S4**). The images were split into 10 regions of interest (ROI) along the length of the microvessel. Heatmaps were comprised of 90 instantaneous permeabilities of 500 kDa dextran at different locations along the microvessel length (ROI 1-10, x-axis) and in ten-minute increments (y-axis). These results show that: (1) melittin-induced BBBO is heterogenous in space and time, and (2) the extent and duration of BBBO is dose dependent.

**Fig. 3.**
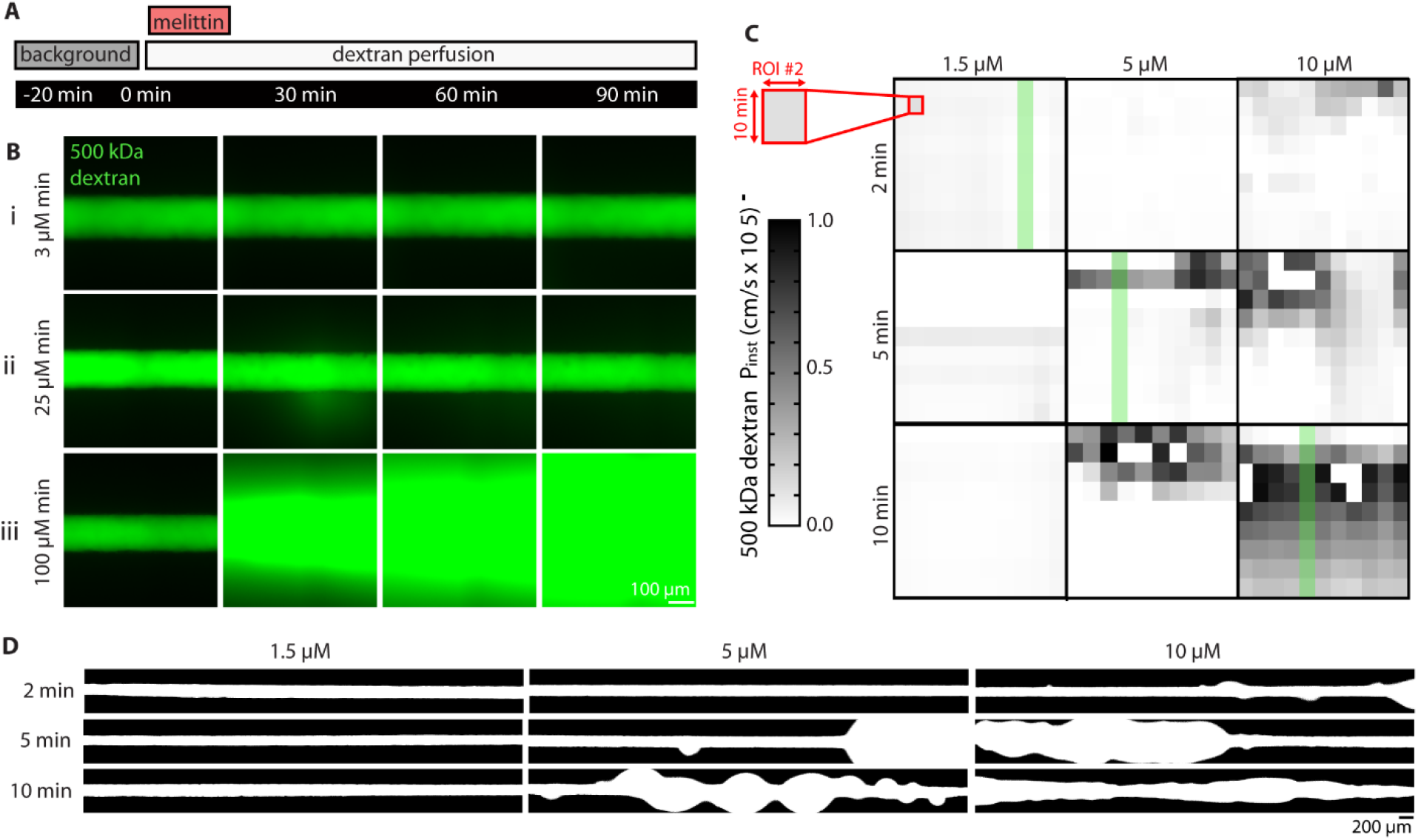
Melittin-induced BBB opening within tissue-engineered dhBMECs microvessels. **(A)** Experimental imaging protocol: microvessels were exposed to 1.5, 5 or 10 μM melittin for two, five, or ten minutes. During and after melittin exposure microvessels were perfused with 500 kDa dextran for 90 minutes. **(B)** Time course fluorescence images for a 2-minute x 1.5 μM (low) dose, a 5-minute x 5 μM (intermediate) dose, and a 10-minute x 10 μM (high) dose of melittin. **(C)** Heatmaps of instantaneous 500 kDa dextran permeability for each melittin dose: large squares are comprised of 90 values representing instantaneous permeability across ten regions of interest (ROIs) along the microvessel length (x-axis) and 9 ten-minute windows (y-axis). The green bar corresponds to the ROI displayed in Fig. 3A. **(D)** Focal leaks visible from binarized images of 500 kDa dextran fluorescence at 30 minutes after initial melittin exposure for each dose. Data collected across *n* = 9 microvessels exposed to doses from 3 – 100 μM·min.

Intermediate and large doses of melittin resulted in the formation of punctate focal leaks, similar to those observed during continuous melittin exposure (**Figs. 3B and S5**). Imaging revealed that the threshold dose for inducing focal leaks was 20 μM·min (**Fig. 3D**). At intermediate doses (20 – 50 μM·min), focal leaks were transient indicating highly reversible BBBO (**Supp. Video 3**).

The average permeability of 500 kDa dextran in melittin-exposed microvessels was dose-dependent (**Fig. 4A**). There was a strong positive linear relationship between exposure time and average permeability (*r*^2^ = 0.91, *p* < 0.001 significantly non-zero slope). For example, 50 μM·min doses increased the average permeability of 500 kDa dextran in a 200 μm segment to approximately 1.5 × 10^−6^ cm s^−1^, ~25-fold higher than baseline. Permeability was simultaneously measured for 3 kDa, 10 kDa and 500 kDa dextran, where all three dyes displayed similar leakage dynamics with higher permeability values at lower molecular weights (data not shown). Additionally, the focal leak density was dose dependent (**Fig. 4B**) (*r*^2^ = 0.58, *p* = 0.017 significantly non-zero slope). At intermediate melittin doses there was a linear relationship between the dose and focal leak density, however, at high melittin doses a similar focal leak density was observed. The linear increase in dextran permeability and the plateau in focal leak density suggested that BBBO is not only related to the number of disruption sites but also the size and duration of opening at each site. Focal leaks typically emerged immediately after melittin administration; however, at longer exposure times focal leaks were also observed during dosing.

**Fig. 4.**
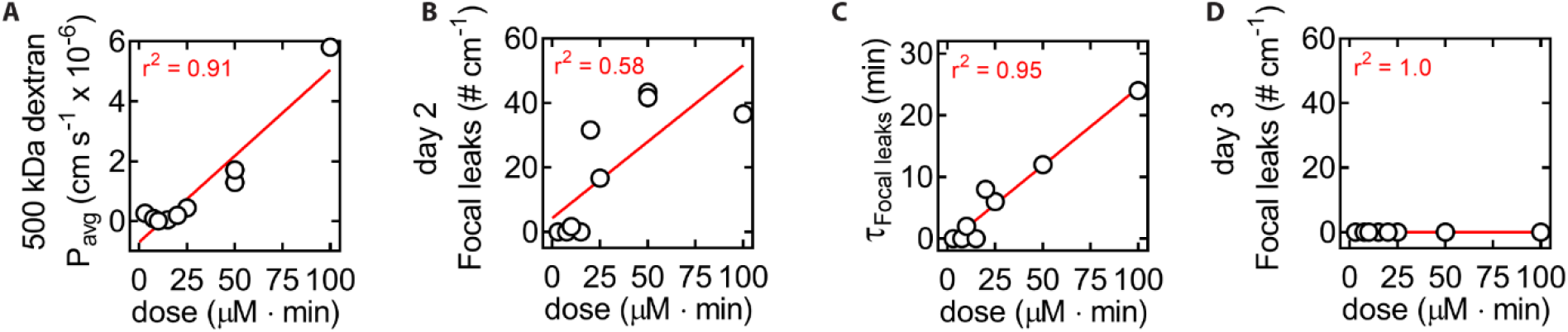
Dose dependence of melittin-induced BBB opening within tissue-engineered dhBMECs microvessels. The melittin dose (in μM·min) is defined as the product of melittin concentration and melittin perfusion time. **(A)** Average permeability (*P*_avg_) of 500 kDa dextran. **(B)**500 kDa dextran cumulative focal leak density. **(C)**500 kDa dextran focal leak reversibility constant, defined as the time at which the last new focal leak emerged after melittin dosing. **(D)** Focal leak density 24 hours after dosing . Data collected across *n* = 9 microvessel exposed to doses from 3 – 100 μM·min. Red lines represent lines of best fit.

The duration of BBBO is an important metric to guide drug delivery strategies since it provides a window of opportunity for enhancing entry into the brain. As a metric for the duration of opening we identified the time at which the last focal leak appeared, which is defined as the focal leak reversibility time constant (τ_focal leaks_). There was a strong linear relationship between melittin dose and τ_focal leaks_ (*r* = 0.98, *p* < 0.001 significantly non-zero slope) (**Fig. 4C**). Additionally, for clinical use, BBBO must be reversible with recovery of normal BBB function after drug delivery. We have previously shown that dhBMEC microvessels display stable permeability over six days with no focal leaks observed [29]. To confirm that melittin-induced BBBO was fully reversible, microvessels were perfused with 500 kDa dextran 24 hours after melittin exposure. Focal leaks were not observed for all melittin doses (3 – 100 μM·min range) indicating recovery of barrier function (**Figs. 4D and S6**).

### 3.4. Melittin variants also support reversible BBBO and highlight possible structure-function relationships required for activity

To determine whether the effects of melittin on BBBO are sequence-specific, we characterized the activity of a scrambled version of melittin (Mel-Scramble) (**Supp. Table 1 and Fig. S7**). The melittin structure contains a high number of cationic amino acids (R, K) at the carboxy terminus contributing to its primary amphipathicity. In addition, melittin is largely alpha helical with hydrophobic and charged amino acids separated on different faces, giving it secondary amphipathicity. We redistributed the amino acids in Mel-Scramble to have no primary amphipathicity and minimal secondary amphipathicity. Circular dichroism measurements showed that Mel-Scramble is partially alpha helical, and the helicity is lower than that of melittin (**Fig. S8**). Mel-Scramble did not induce BBBO for 25, 50, or 100 μM·min doses. There was no change in the permeability of 500 kDa dextran following perfusion with Mel-Scramble and no focal leaks were observed during or 24 hours after dosing (**Fig. S9**). There was a non-statistically significant relationship between dose and 500 kDa dextran permeability (*p* = 0.335), focal leak density (*p* = not calculable), and focal leak reversibility (*p* = not calculable) (**Fig. S9 A-D**).

Having established that the action of melittin is sequence-specific, we assessed the effects of small sequence modifications on BBBO. Specifically, we assessed the ability of melittin sequence variants that are known to be membrane active (i.e. Melittin-CONH_2_, MelP3, and MelP5) to induce reversible BBBO in microvessels (**Supp. Table 1 and Fig. S8**). Thus far, all BBBO experiments were performed using melittin with a carboxy terminus (COO^−^). The melittin-CONH_2_ variant is a natural form of melittin in bee venom with a carboxyamide (CONH_2_) terminus. At physiological pH, the carboxyamide is neutral and can facilitate insertion into lipid bilayers, contributing to enhanced membrane binding compared to the carboxy terminus variant. MelP3 and MelP5 were originally discovered in a high-throughput *in vitro* screen for their potent ability to form pores in lipid membranes [26]. We observed that these three melittin variants at a concentration of 10 μM and at 2-, 5- and 10-minute exposures induced reversible BBBO (**Fig. S9E**). Across the closely related four variants, there were non-statistically significant differences in the dose-dependence of 500 kDa dextran permeability (*p* = 0.051) and focal leak density (*p* = 0.415) (**Fig. S9E, F**). However, the reversibility of focal leaks was variant-dependent as there were statistically significant differences in the slope of focal leak reversibility versus dose (*p* < 0.001) (**Fig. S9G**). Specifically, melittin-CONH_2_ displayed the highest slope (**Fig. S9G**), suggesting that disruptions in barrier function may occur over longer time periods than for melittin-COO^−^, despite lower dextran permeability (**Fig. S9E**). We hypothesize that the neutral C-terminus of melittin-CONH_2_ has greater binding to dhBMEC membranes, resulting in extended barrier disruption. Across all variants, focal leaks were not observed 24 hours after dosing showing that barrier disruption was reversible (**Fig. S9H**).

### 3.5. Transient melittin dosing decreases cell turnover

To explore the mechanisms of melittin-induced BBBO, we quantified dhBMEC behavior in microvessels using live-cell imaging (**Fig. 5A**). At melittin doses below 20 μM·min, non-disruptive events including cell division and apoptosis were observed. These events were not associated with loss of barrier function or formation of focal leaks under baseline conditions as surrounding endothelial cells are able to dynamically retain cell-cell contacts [29]. However, at melittin doses above 20 μM·min, junctional disruption was observed resulting in the formation of focal leaks.

**Fig. 5.**
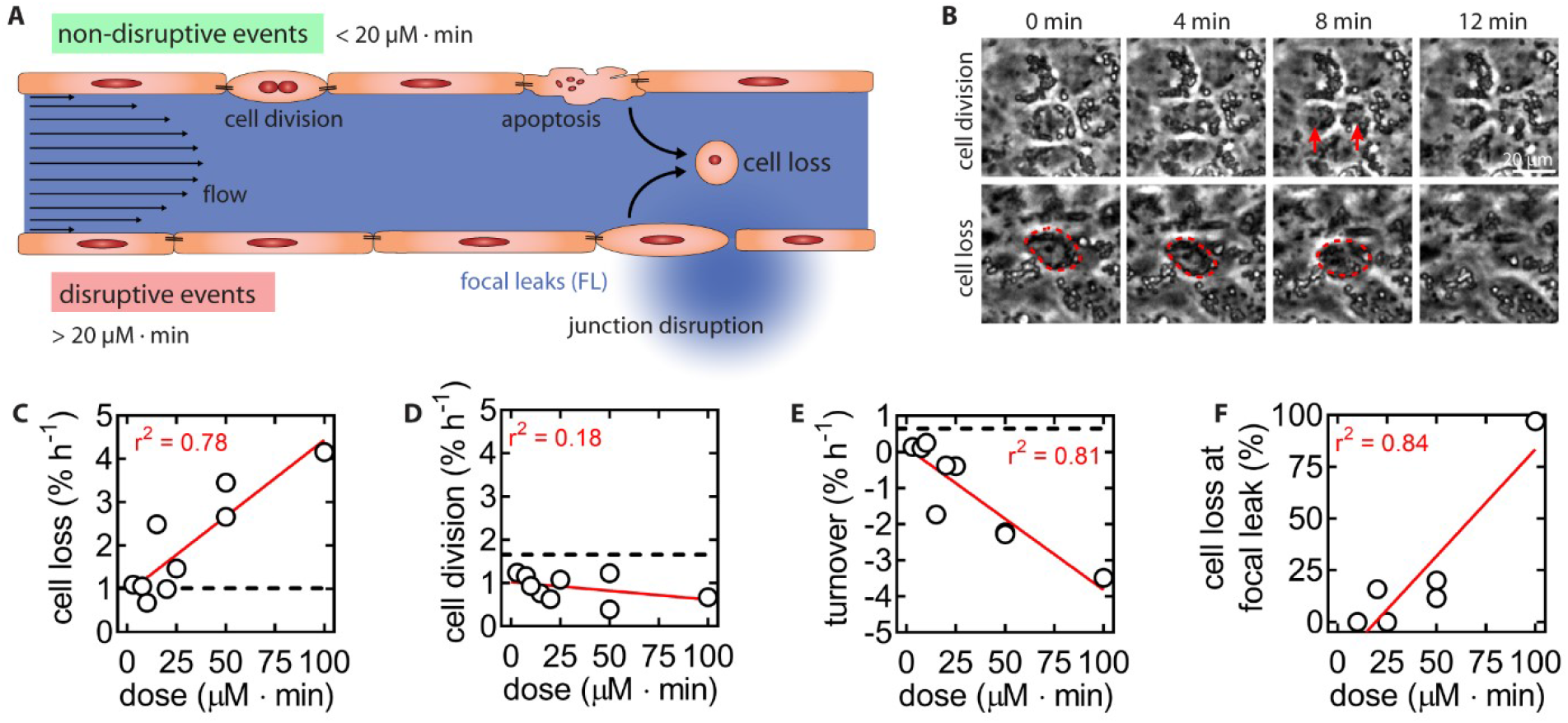
Cellular dynamics of melittin-induced BBB opening within tissue-engineered dhBMECs microvessels. **(A)** Schematic illustration of barrier-disruptive and non-disruptive events observed during melittin exposure. Mitosis and apoptosis (non-disruptive events) do not result in local permeability increases, while junction disruption causes focal leaks of 500 kDa dextran. **(B)** Representative phase contrast time course images of dhBMEC cell loss and cell division in microvessels. Events were manually traced using live-cell phase contrast imaging of dhBMEC monolayers. Mitosis is visible as cell compression, alignment of chromosomes, and the formation of daughter cells; cell loss is visible as a cell shrinkage and subsequent disappearance from the monolayer. **(C-E)** Cell loss, cell division, and net turnover rates for dhBMECs across melittin doses. Cell loss displays a strong positive correlation to melittin dose, while proliferation displays a weak negative correlation. **(F)** Spatial correlation of cell loss events and focal leaks. Percent of focal leaks with detected cell loss within 50 μm of the leak and within 4 minutes of the leak formation versus melittin dose. Data collected across *n* = 9 microvessels exposed to doses from 3 – 100 μM·min. Red lines represent lines of best fit. Black dotted lines represent mean rates for dhBMEC microvessels under homeostatic conditions.

To evaluate the effect of melittin dosing on dhBMEC behavior, we manually traced cell loss and cell division events in phase contrast images (**Fig. 5B**). Under baseline conditions, the loss of cells from dhBMEC microvessels was previously found to be ~1% h^−1^ [29]. The cell loss rate following melittin exposure was dose-dependent (*r*^2^ = 0.78, *p* = 0.002 significantly non-zero slope) (**Fig. 5C**). However, we did not detect focal leaks when the cell loss rates were similar to baseline conditions, showing that apoptosis can occur without disruptions in barrier function, as described above. The cell division rate was about 1% h^−1^ but was not melittin dose-dependent (*r*^2^ = 0.18, *p* = 0.260 not significantly non-zero slope) (**Fig. 5D**), but was lower than under control conditions (~1.7% h^−1^) [29], suggesting that melittin inhibits proliferation across tested melittin doses. The turnover of dhBMECs in microvessels, defined as the cell division rate minus the cell loss rate, was negative and displayed a strong dose dependence (*r*^2^ = 0.81, *p* < 0.001 significantly non-zero slope) (**Fig. 5E**). Under baseline conditions, BBB microvessels displayed a turnover of ~0.6 % h^−1^, suggesting a net gain of cell density over time [29]. The strong correlation between cell turnover and dose suggests that local cell loss is responsible for BBBO. Interestingly, the loss of cells from microvessels was spatially correlated with focal leaks (*r*^2^ = 0.84), further suggesting a shared mechanism (**Fig. 5F**).

### 3.6. Transient melittin dosing increases paracellular permeability via disruption of cell junctions

We further investigated the mechanism of melittin-induced BBBO using epifluorescence and confocal microscopy. Epifluorescence images at the microvessel mid-plane showed that melittin exposure resulted in cell swelling (**Fig. 6A**). Dextran exclusion from the endothelium suggested that leakage was due to disruption of paracellular barrier function. This was confirmed using confocal microscopy (**Fig. 6B**). Disruption was spatially localized since the endothelium was intact above and below each site, suggesting that focal leaks originated from discrete disruptions of cell-cell junctions. To further assess this hypothesis, we performed immunohistochemistry for the tight junction (TJ) scaffolding protein zona occludens-1 (ZO1) after melittin treatment of 2D dhBMEC monolayers across doses from 3 μM·min to 450 μM·min (matching the doses used in the cell viability assay). Melittin doses of 3 – 100 μM·min, within the range of reversible BBBO, did not result in large changes in ZO1 protein expression compared to controls (**Fig. S10**). However, above the limit of reversibility at a 450 μM·min dose significant cell detachment was observed, similar to the results from the viability assay and microvessels at high melittin doses (**Figs. S2 and S5**). Interestingly, cells that bordered the defects in the monolayer frequently showed ZO1 staining (**Fig. S10**; marked with red arrows). Thus, we hypothesized that BBB disruption occurs by mechanisms related to local disruption of TJs rather than a global decrease in TJ integrity.

**Fig. 6.**
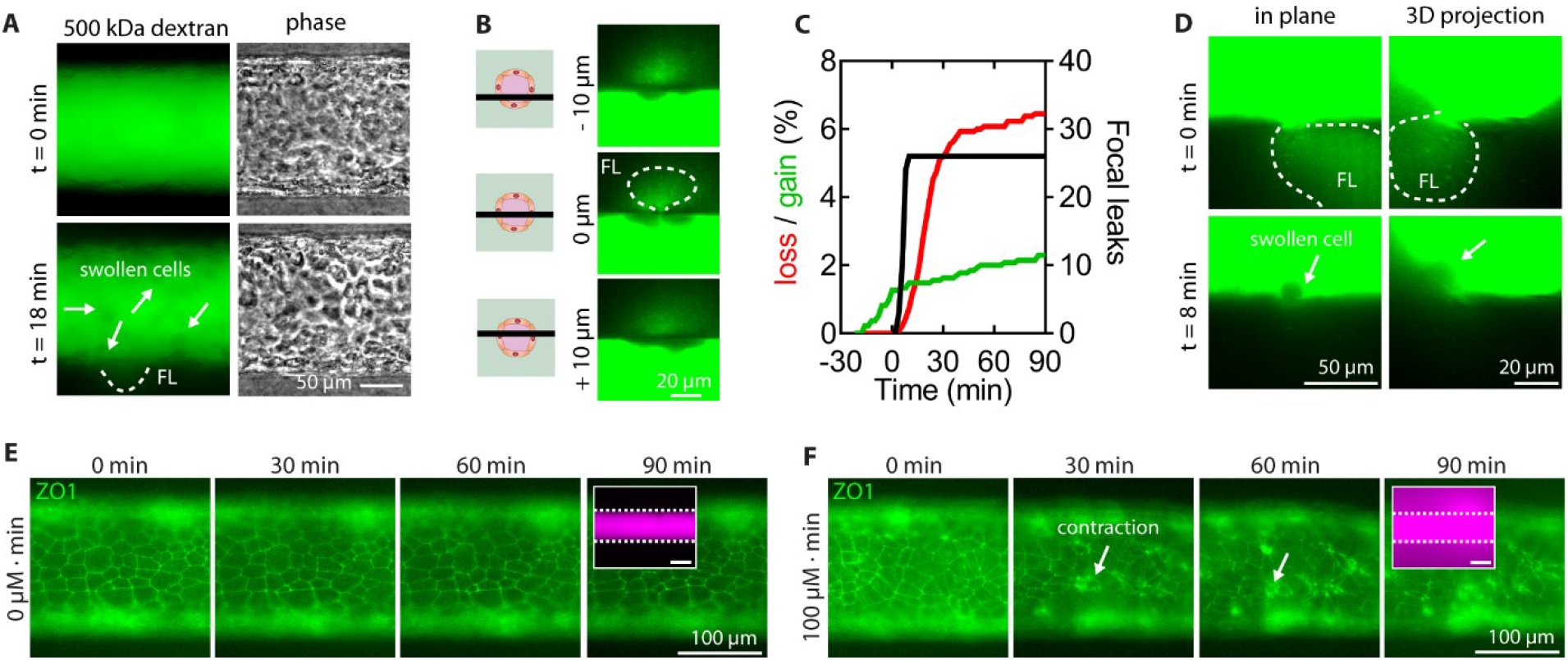
Mechanisms of junctional disruption during melittin-induced BBBO within tissue-engineered dhBMEC microvessels. **(A)** Epifluorescence images of 500 kDa dextran-fluorescein at 0 min and 18 minutes (8 minutes after a ten-minute 10 μM melittin perfusion). Swollen cells visible due to dextran exclusion (white arrows). **(B)** Focal leaks are visible as paracellular plumes, and are restricted to local disruption of cell-cell junctions: 10 μm above or below this location the endothelium is intact. The schematic illustrations show the cross-sectional locations of the confocal images. **(C)** Time course of cumulative cell loss, cell gain, and focal leak formation for a 50 μM·min melittin dose (other doses shown in supplemental information). Event counting was initiated 30 minutes before dosing. **(D)** Cell swelling occurs at the site of focal leaks after focal leak formation. **(E)** Tight junctions display stable localization under baseline conditions in dhBMECS with GFP-tagged ZO1. **(F)** Exposure to melittin induces cell contraction (white arrows) of discrete dhBMECs. Confocal images were collected across *n* = 3 BBB microvessels exposed to 50 μM·min melittin.

To better understand if focal leak formation during melittin exposure was associated with cell loss, we plotted the cumulative number of cell loss events, cell division events, and focal leaks over two hours following dosing in microvessels (**Fig. S11**). For a representative 50 μM·min dose, the number of cell loss events increased over background after a plateau in the number of focal leaks (**Fig. 6C**). Across other melittin doses, increases in cell loss was consistently observed after focal leak formation (**Fig. S11**). Thus, the removal of a cell from the endothelium was not responsible for focal leak formation. Instead, time-course confocal imaging showed that focal leaks emerged before cell swelling and with high spatial confinement (**Fig. 6D**).

To understand the process that leads to the formation of focal leaks, we performed additional experiments on microvessels comprised of dhBMECs with GFP-tagged zona occludens-1 (ZO1) [43]. This cell line allowed us to visualize tight junction localization during real-time imaging. Under baseline conditions (no melittin), tight junctions maintained stable localization over 90 minutes with no leakage of 10 kDa dextran (**Fig. 6E**). However, upon exposure to high melittin doses (100 μM·min), cells contracted, as evident by condensation of ZO1 at discrete locations in the monolayer (**Fig. 6F**, **Supp. Video 4**). These results suggest that melittin induces cell contraction which transiently disrupts cell-cell junctions and results in the formation of focal leaks. We observed that cells that swelled into the microvessel lumen following contraction were eventually removed by the shear flow. These results are consistent with the observed spatial correlation between the loss of cells and focal leaks (*r*^2^ = 0.84) (**Fig. 5F**).

### 3.7. Profile of melittin-induced BBBO in a mouse model

To assess BBBO *in vivo*, melittin was infused into an internal carotid artery in a mouse model following catheterization. The internal carotid artery predominately supplies the ipsilateral hemisphere of the mouse brain, while other catherization protocols could be used to distribute BBB downstream of any major human artery. In preliminary experiments, we studied three intra-arterial (IA) administrations: 1, 3, or 5 μM melittin for 1 minute (1 – 5 μM·min). To determine the minimum effective dose, we perfused Evans Blue immediately following the injection to determine BBB status. 3 μM·min melittin was identified as the minimum effective dose by reproducible leakage of Evans Blue within the downstream ipsilateral hemisphere following injection (**Fig. S12**). No leakage of Evans Blue was observed following 1 μM·min melittin, indicating that IA administration itself does not compromise BBB status as previously found [44, 48]. Widespread leakage of Evans Blue was observed following a 5 μM·min melittin dose, however, mice displayed rotational behavior indicating neurological damage to one hemisphere of the cortex. Thus, 3 μM·min melittin was further explored for efficacy and safety using MRI and histopathology.

Utilizing dynamic T1-weighted MRI, we found that BBBO occurred during injection of 3 μM·min melittin, indicating nearly instantaneous disruption of brain capillaries (**Supp. Video 5**). Leakage of Gadolinium (Gd) and Evans Blue was observed immediately following dosing, and normal barrier function was confirmed 24 hours and 7 days later (**Fig. 7A, B**). Following dosing there was a significant increase in T1 intensity in the ipsilateral hippocampus and thalamus/hypothalamus compared to the contralateral counterpart (*p* = 0.021 and 0.033, respectively) (**Fig. 7C**). T1 hyperintensity in the cortex was much lower indicating limited BBBO (*p* = 0.134). Follow-up T1 MRI showed marginal leakage of Gd at 24 hours (*p* = 0.154) and no Evans Blue extravasation, indicating rapid recovery of barrier function. On day 7 both Gd-enhanced MRI (*p* = 0.169) and Evans Blue confirmed full recovery of the BBB after 3 μM·min melittin exposure. These observations were not brain-region specific; no statistically significant changes in T1 intensity were found in any brain regions at 24 hours and 7 days (*p* > 0.05). Additionally, using thresholding of T1 intensity we found that a 3 μM·min dose opened a brain territory of 5.70 ± 1.73 mm^2^ within the hippocampus, which reverses to ~20-fold less in area at 24 hours and 7 days (*p* = 0.015 and 0.035, respectively) (**Fig. 7D**).

**Fig. 7.**
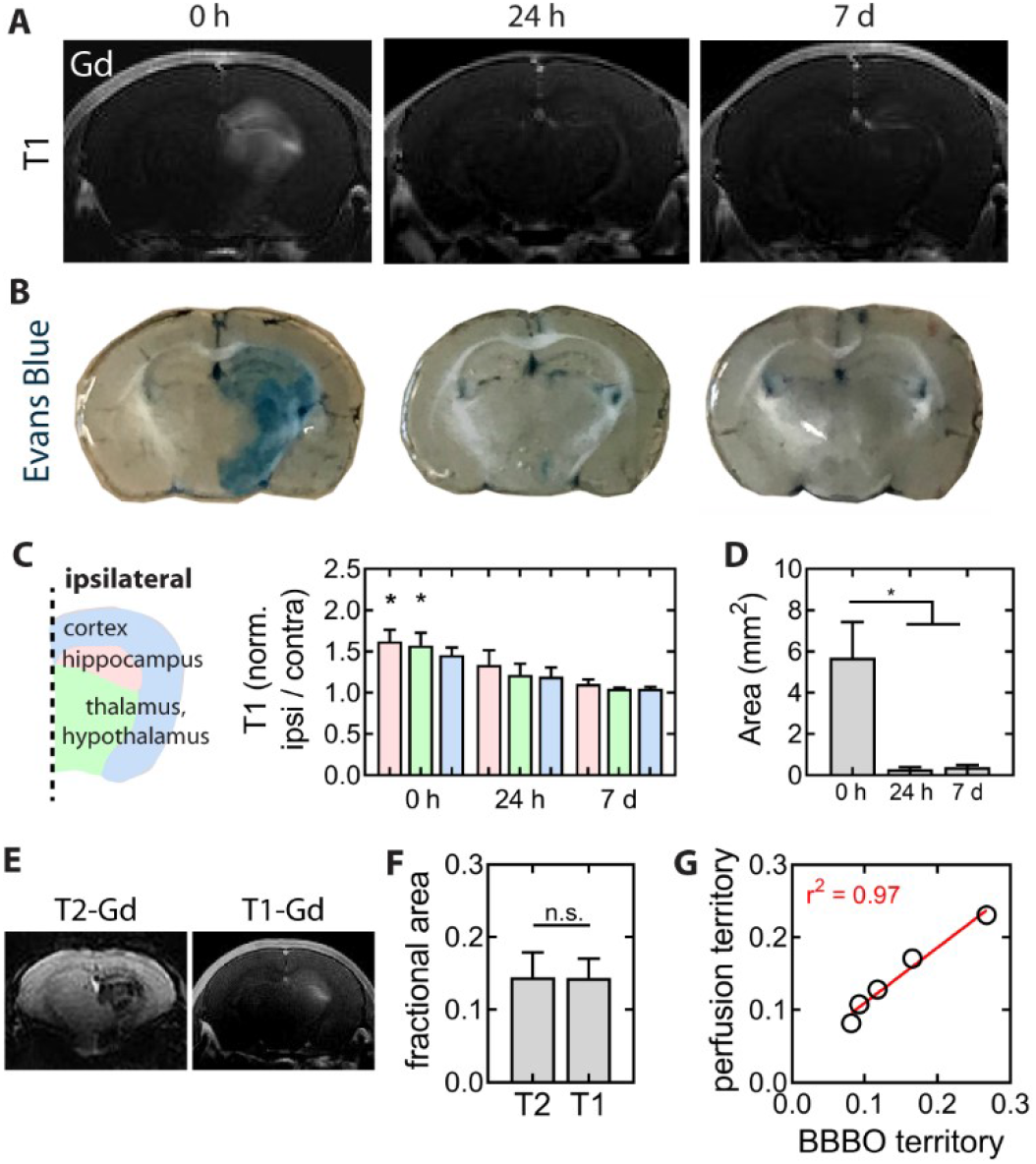
Profile of melittin-induced BBBO in mice. **(A-B)** Representative time course of T1-weighted gadolinium-enhanced magnetic resonance imaging and Evans Blue leakage following 3 μM·min melittin-induced BBB opening. **(C)** Brain-region specific dynamics of melittin-induced BBBO. A significant difference between T1 intensities in the ipsilateral and contralateral hemispheres is only observed in the hippocampus and thalamus/hypothalamus immediately after melittin dosing. **(D)** Quantification of the area of BBBO using thresholding of T1 hyperintensities. *n* = 4, 5 and 3 mice at 0 h, 24 h and 7 days, respectively for the 3 μM·min melittin dose. **(E)** T2-weighted gadolinium-enhanced imaging was conducted prior to BBBO to correlate perfusion territories and BBBO territories. **(F-G)** The fractional areas of brain perfusion and BBBO are well correlated (*n* = 5). * p < 0.05. Bars represent mean ± SEM.

While 3 μM·min melittin consistently induced BBBO, there was variability in the extent of opening, ranging from small regions, limited to the hippocampus, to large areas encompassing the majority of the ipsilateral hemisphere (**Fig. S13A**). However, this variability was not melittin-specific, as hyperosmotic approaches utilizing injection of 1.4 M mannitol for 1 minute displayed similar variability patterns (**Fig. S13B**). Additionally, the extent of BBBO was similar between both approaches as measured by similar territories of opening (p = 0.333) (**Fig. S13C**). This variability stems from the highly integrated and collateralized cerebral circulation from the four major cerebral arteries and microregulation dynamically routing cerebral blood flow. This is a well-established phenomenon and a major driver of variability in clinical outcomes with IA drug delivery into the brain.

To address this challenge and improve the predictability of the territory of BBBO we have developed a technique based on interventional MRI where injection of the BBB-opening agent is preceded by dynamic contrast-enhanced MRI during IA infusion of a contrast agent (**Fig. 7D**), as previously described [48, 49]. Intra-arterial infusion of saline supplemented with 50 mM gadolinium with dynamic T2-weighted MR imaging facilitates detection of hypointense signal from the brain region selectively perfused by the microcatheter. The infusion rate can be manipulated to achieve effective perfusion of a desired brain territory and at that point melittin is infused intra-arterially to the same brain region. With that approach we were able to predict the BBBO territory as there is excellent agreement between contrast-perfusion territory and BBBO territory (**Figs. 7E,F and S14**) (*r*^2^ = 0.97, *p* = 0.002 significantly non-zero slope).

### 3.8. Safety of melittin-induced BBBO

In 11 mice treated with 3 μM·min melittin, all animals except one woke up from anesthesia without apparent neurological deficits (paralysis, seizure, or rotation). One mouse in this group died within 12 h, possibly due to complications of surgery. In the remaining mice there was no evidence of neurological deficits and the cohort dedicated for long-term survival (30 days) reached that endpoint without complications.

The effects of melittin-induced BBBO on edema and inflammation were studied for a 3 μM·min dose at acute (24 hours) and chronic (7 days) time points. Follow-up MRI at 24 h following a 3 μM·min dose showed no evidence of edema or inflammation on T2 scans or micro-hemorrhages on susceptibility weighted imaging (SWI) scans (**Fig. S15A**). Similarly, no abnormalities were detected in MRI at 7 days (**Fig. 8A**), providing compelling evidence of neurological safety. Since the hippocampus was the region with the most consistent BBBO, we focused on that region for histopathological assessment. Staining for eriochrome cyanine revealed normal myelin at 7 days (**Fig. 8B**). The activation of astrocytes and microglia was assessed from immunohistochemistry images of glial fibrillary acidic protein (GFAP) and ionized calcium binding adaptor molecule 1 (Iba1) (**Fig. 8C**). Both sensitive markers of neuroinflammation showed no statistically significant differences in intensity between the ipsilateral and contralateral hemispheres 7 days after treatment (*p* = 0.628 and 0.372, respectively) (**Fig. 8E**). Neuronal damage was assessed from immunohistochemistry images of the neuron-specific marker NeuN and the apoptotic marker caspase-3 (**Fig. 8D**). There was no statistically significant change in the intensity of both markers in the ipsilateral hemisphere 7 days after treatment (*p* = 0.933 and 0.562, respectively) (**Fig. 8E**). We also evaluated neuroinflammation and neuronal damage at the acute stage, 24 h after IA injection (**Fig. S15B-D**). Both T2 and SWI appeared normal, and GFAP, Iba1, NeuN, and caspase-3 intensities were not statistically different between hemispheres (*p* > 0.05 for all cases).

**Fig. 8.**
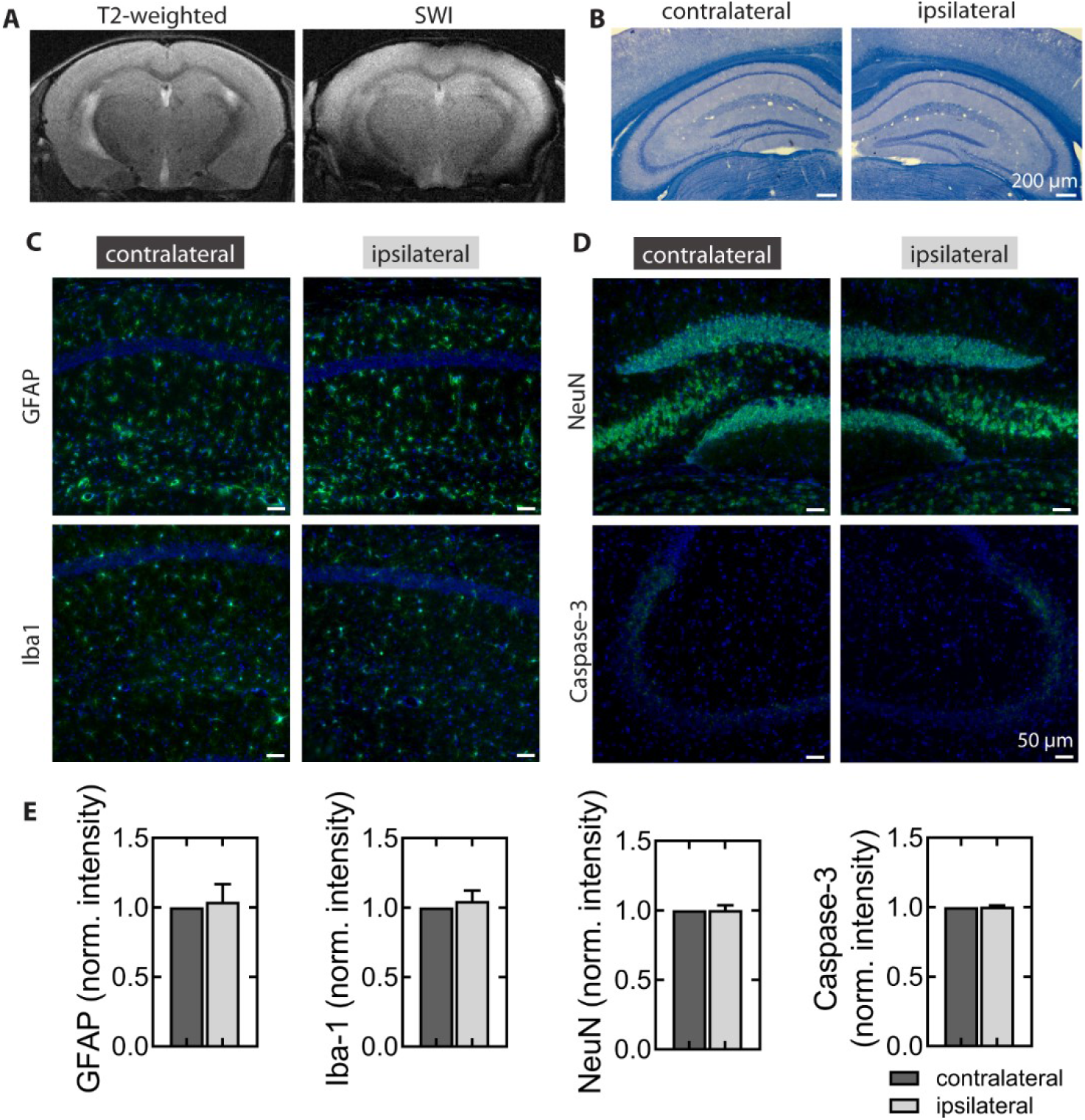
MRI and histological assessment 7 days following melittin-induced BBB opening at a dose of 3 μM·min. **(A)** T2-weighted and SWI images 7 days after BBBO showed no sign of brain damage or microhemorrhage, respectively. **(B)** Histological staining with eriochrome cyanine revealed that there was no myelin demyelination 7 days after BBBO. **(C)** Immunohistochemical detection of neuroinflammation markers glial fibrillary acidic protein (GFAP) and ionized calcium binding adaptor molecule 1 (Iba1) in the contralateral and ipsilateral BBBO region. **(D)** Immunohistochemical detection of damage markers using a neuronal nuclear antigen (NeuN) and apoptotic marker (caspase-3) in the contralateral and ipsilateral BBBO region. **(E)** Comparison of normalized intensity of immunohistochemical images between the contralateral (control) and ipsilateral BBBO region. Representative images are shown. *n* = 3 mice for each imaging technique or stain. Bars represent mean ± SEM.

## 4. Discussion

### 4.1. Reversible BBB opening by optimizing melittin dose

The BBB is a major roadblock for drug, gene, and cell delivery to the brain. We demonstrate that when optimally dosed, melittin induces reversible BBB opening (BBBO) via enhanced paracellular permeability in a tissue-engineered BBB microvessel model. Other melittin sequence variants (MelP3 and MelP5) induce reversible BBBO, however, a scrambled melittin sequence (Mel-Scramble) did not induce BBBO, demonstrating sequence specificity. When administered as an intra-arterial injection into the mouse cerebrovasculature, melittin supports reversible BBBO without neurological damage. Importantly, intra-arterial MAP-induced BBBO has distinct advantages over existing approaches, including: (1) dramatic fold-increases in brain penetration for a wide range of molecular weights, (2) opening far below the solubility limit, (3) a large peptide design space to optimize opening, reversibility and therapeutic effect, (4) avoiding systemic injections to achieve BBBO, and (5) a lack of systemic and neuronal toxicity due to intra-arterial doses which are diluted upon exiting the brain. Existing techniques to modulate BBB permeability result in only modest increases in therapeutic penetration or are highly inefficient. For example, efflux pump inhibition results in a 2-3 fold increase in efflux substrate permeability [4, 29], while MR-guided focused ultrasound (FUS) is highly inefficient as only a small fraction of systemically administered therapeutics enter the brain (as low as 0.009% of injected dose) [50]. Intra-arterial melittin is particularly suited for delivery of antibodies (MW ~150 kDa) which display negligible penetration into the healthy brain.

### 4.2. Comparison of melittin and mannitol

Comparison of melittin-induced and hyperosmotic BBBO *in vivo* showed that the dynamics were similar, however, melittin-induced BBBO occurred well below the solubility limit of melittin. Intra-arterial injection with 3 μM·min melittin (~ 120-fold below solubility limit) resulted in robust opening of the mouse hippocampus and thalamus/hypothalamus. This suggests that BBBO could be implemented without complete displacement of blood volume as required for mannitol which only induces opening close to its solubility limit [9, 10]. Additionally, we conducted retrospective analysis to compare mannitol [10] and melittin BBBO within tissue-engineered BBB microvessels (**Fig. S16**). For both agents, a critical threshold dose must be exceeded to induce opening; for mannitol, this threshold was 7 M·min, while for melittin it was 20 μM·min. Melittin may be able to alter BBB permeability more potently, as above these thresholds melittin resulted in a 3-fold higher focal leak density.

The variability in opening territory in the mouse brain following melittin-induced BBBO is also observed using hyperosmotic BBBO, and is likely due to differences in blood supply and collateral flow [51]. To address this variability, we utilized real-time imaging of the catheter perfusion territory to monitor BBBO. This is a critical step in translation towards clinical use [52]. Previously, we have shown with interventional dynamic susceptibility contrast-enhanced MRI that the local trans-catheter perfusion territory is indeed highly variable [48]. In addition to visualizing trans-catheter perfusion, interventional MRI could be exploited to improve the precision and predictability of the BBB opening territory [49].

### 4.3 Safety of the technique

The safety of BBBO is critical for clinical applications. MAP-based therapies have been utilized for treatment of cancer, diabetes, inflammation, atherosclerosis, arthritis, epilepsy and neurodegeneration in animal models [53, 54]. However, challenges include non-specific cytotoxicity, peptide degradation, and hemolytic activity [55]. In mice, the LD50 dose for intraperitoneal administration of melittin is 3.2 mg kg^−1^ [56]. While the regional concentration of melittin in the brain is high after IA administration, the systemic dose is ~ 50-fold lower than the LD50 (0.64 mg kg^−1^ for 3 μM·min melittin, assuming a mouse weight of 20 g). Additionally, assuming dilution in the bloodstream upon exiting the brain, melittin concentrations would be reduced ~10-fold (0.33 μM for 3 μM·min melittin, assuming a 150 μL injection is diluted into 1.2 mL of whole blood). Thus, by utilizing IA delivery into the carotid artery we minimize systemic toxicity.

Although the majority of enhanced permeability into the brain occurs after the melittin dose (as observed at intermediate doses in tissue-engineered microvessels), BBB disruption during dosing may result in melittin leakage from circulation into the brain parenchyma. In our tissue-engineered BBB model we observed transport of 500 kDa dextran into the ECM during melittin perfusion, which implies that melittin (MW 2.7 kDa), can also be transported into the surrounding tissue during the perfusion. In our cytotoxicity assay we showed that HCNs maintained high viability (≥ 98%) for ~12 minutes in 5 μM and 10 μM melittin, and ~23 minutes for 0.75 μM melittin. For *in vivo* experiments we used a 1-minute dose and hence neuronal exposure to melittin in the vicinity of focal leaks is expected to be short. Therefore, we speculate that melittin will not cause significant neurotoxicity even at relatively high local doses. This conclusion was further supported by *in* vivo toxicity studies, finding limited neurotoxicity, demyelination or neuroinflammation after seven days or neurological deficits for up to 30 days.

The safe dose range for intra-arterial injection is relatively narrow in mice since 1 μM·min melittin did not result in BBBO and 5 μM·min melittin resulted in neurological deficits. A narrow regime of neurological safety is also observed in mice for hyperosmotic BBBO, which requires high injection speeds to achieve complete blood displacement. Excessive infusion rates can be damaging, leading to brain lesions of microembolic appearance [57]. The 5 μM·min melittin dose induced substantial BBBO, with enhancement throughout most of the ipsilateral hemisphere as visualized using Evans Blue leakage. Across 7 mice, neurological deficits were apparent after the procedure, which worsened over time. Specifically, rotational mobility was observed suggesting damage to the ipsilateral cortex. Thus, damage in response to the 5 μM·min melittin dose was likely due to neuronal toxicity due to dramatic BBBO.

### 4.4 Mechanisms of melittin-induced BBB opening

Utilizing our tissue-engineered microvessels we identified the key mechanistic features of intra-arterial BBBO: (1) increased paracellular (but not transcellular) permeability, (2) focal leaks associated with cell contraction, and (3) reversible junction disruption based on dose. Melittin does not globally change the localization or expression of tight junctions (as observed using immunocytochemical detection of zona occludens-1). Instead, we observe that melittin induces local cell contraction. This matches previous studies which find that melittin causes calcium influx due to pore formation in cell membranes [58, 59]. Calcium flux into the endothelium is a key regulator of cytoskeletal organization and barrier function, and has been shown to induce disruption of tight junctions [60, 61]; other chemical BBBO agents are reported to function via a similar mechanism [19, 62]. Therefore, we suggest that cell contraction induces strain on surrounding cells that, if sufficiently large, transiently disrupts cell-cell junctions resulting in a focal leak (**Fig. S17**). At low doses (below 20 μM·min in microvessels), the tensile forces on cell-cell junctions are insufficient to induce focal leak formation and the monolayer is able to maintain integrity. However, at intermediate doses (20 μM·min – 50 μM·min) the tensile forces resulting from cell contraction are sufficiently large to induce local disruption of weak cell junctions resulting in the formation of focal leaks. The density of contraction and disruption events is sufficiently low that they are isolated, and recovery of normal barrier function occurs rapidly. At high doses (100 μM·min) the density of contraction events is sufficiently high that clusters of disrupted cell-cell junctions can result in cell loss; however, this regime is not observed *in vivo* as hemorrhaging does not occur.

### 4.5 In vitro microvessels as a predictive tool for in vivo BBBO

We identified an approximately 7-fold difference in melittin dose required for BBBO in tissue engineered microvessels (20 μM·min) compared to the mouse brain (3 μM·min). Assuming that our tissue-engineered model accurately mimics the human BBB, then this difference could arise from species-to-species differences in barrier function [63]. Support for this explanation comes from the fact that other enhancers of brain permeability appear to display a dose mismatch between rodents and humans [64, 65]. However, since there are insufficient data on permeability or other quantitative measures of barrier function in the human brain, we cannot confirm that our tissue-engineered model accurately mimics barrier function in the human brain. We have previously found that parameters including diameter, shear stress, matrix mechanics, and the presence of pericytes do not significantly alter baseline permeability within tissue-engineered BBB models [66, 67]. However, it remains to be determined if these same parameters alter the response of *in vitro* models to melittin or other BBBO agents.

The findings of this study have important clinical applications for delivery of therapeutics into the brain. We validate the use of tissue-engineered BBB models in developing clinically relevant strategies for drug delivery. Specifically, we demonstrate the potential for MAPs to reversibly open the BBB. Clinical translation will require further *in vitro* and *in vivo* validation. Safety and dose optimization studies need to be conducted in large animals, where IA catheters can be advanced more distally for super-selective targeting similar to that performed in humans. Due to the differences in brain and blood volume between mice and humans, neuronal and systemic toxicity may vary widely, and hence the dosing space should be further explored to identify regimes for robust BBBO with the greatest safety margin. Our ultimate goal is to engineer human BBB models with improved predictive power for clinical translation by studying the effect of model parameters on BBBO and comparing results between *in vitro* and *in vivo* models.

## Data and materials availability

All data associated with this study are available in the main text or the supplementary materials. The raw data required to reproduce these findings are available from the corresponding author on reasonable request.

## Acknowledgements

This work was supported by NIH/NINDS (R01NS106008, R01NS09111, R01NS102675), DTRA (HDTRA1-15-1-0046), and NSF (DMR 1709892). RML acknowledges a National Science Foundation Graduate Research Fellowship under Grant No. DGE1746891, AK acknowledges support from a Kirschstein-NRSA Individual Predoctoral Fellowship (F31) under award number NINDS 1F31NS101875, JGD acknowledges support from the Nanotechnology for Cancer Research training program. We also acknowledge Dr. William Wimley (Tulane University) for useful discussions and materials, and Matt Sklar, Gabrielle Grifno, Alanna Farrell, and Erin Gallagher for assistance with cell culture and microfabrication.

## Author contributions

RML, AK, KH and PCS conceived the study. RML and PCS wrote the paper, with input from AK, XL, PW and KH. RML, AK, and JGD conducted and analyzed *in vitro* experiments. XL, CC, RML and PW conducted and analyzed *in vivo* experiments. GL analyzed *in vivo* experiments related to BBBO territory. All authors reviewed and accepted the manuscript.

