## Supplemental Information for "Reversible blood-brain barrier opening utilizing the membrane active peptide melittin *in vitro* and *in vivo*"

**SUPPLEMENTARY INFORMATION**

**Figures**

Figure S1. Melittin-induced disruption of stem-cell derived brain microvascular endothelial cell-like cells (dhBMECs) in a transwell assay.

Figure S2. Concentration-dependent melittin-induced cytotoxicity on derived human BMEC-like cells (dhBMECs) and human cortical neuron cells (HCNs).

Figure S3. Fabrication and perfusion of tissue-engineered BBB microvessels.

Figure S4. Analysis of BBB opening and recovery within 3D microvessels.

Figure S5. Melittin-induced BBB disruption of in vitro blood-brain barrier microvessels.

Figure S6. Reversibility of BBB opening 24 hours after melittin exposure.

Figure S7. Chemical structures of peptides used in this study.

Figure S8. Scrambled melittin maintains alpha helicity.

Figure S9. Melittin variants but not scrambled melittin induce reversible BBB opening.

Figure S10. Immunofluorescence imaging of zona occludens-1 (ZO1) junctions after melittin exposure.

Figure S11. Time course analysis of dhBMEC behavior.

Figure S12. Evans Blue leakage assay to determine minimum effective dose.

Figure S13. Comparison of melittin and mannitol-induced BBB opening.

Figure S14. Perfusion territory of intraarterial injections.

Figure S15. MRI and histological assessment 24 hours following melittin-induced BBB opening.

Figure S16. Retrospective comparison of mannitol and melittin-induced BBBO within tissue-engineered microvessels.

Figure S17. Mechanisms of melittin-induced BBBO.

**Tables**

Table S1. Amino acid sequences and properties of melittin variants.

Table S2. Primary antibodies used in this study.

**Movies**

Movie S1. BBB microvessels display negligible permeability to 500 kDa dextran.

Movie S2. BBB disruption during continual perfusion of 5 μM melittin.

Movie S3. Reversible BBB opening with a 10 min 5 μM melittin dose.

Movie S4. Imaging transient BBB opening using microvessels comprised of zona-occludens-1 fluorescently-tagged dhBMECs.

Movie S5. Real-time imaging of BBB opening of mouse brain using 3 μM melittin for 1 minute under gadolinium enhancement on T1-weighted imaging.

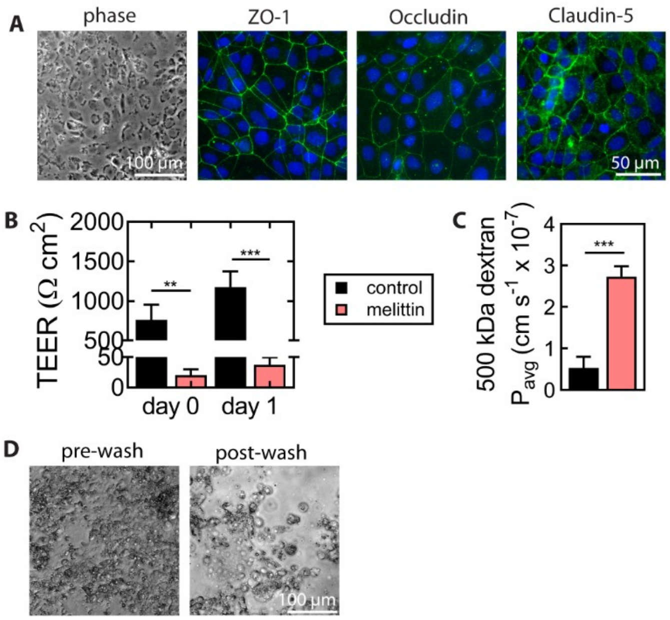


**Fig. S1.** Melittin-induced disruption of stem-cell derived brain microvascular endothelial cell-like cells (dhBMECs) in a transwell assay. **(A)** Under baseline conditions dhBMECs monolayers express tight junction proteins zona occludens-1 (ZO-1), occludin, and claudin-5. **(B)** Transendothelial electrical resistance (TEER) of dhBMEC monolayers before and after exposure to 5 µM melittin for 90 minutes on the day of the exposure (day 0) and the next day. **(C)** 500 kDa dextran permeability across dhBMECs is increased after treatment with 5 µM melittin for 90 minutes. **(D)** Phase contrast image of dhBMECs cultured on glass slides and treated with 5 µM melittin for 90 minutes before and after washes with buffer. Melittin treatment results in cell swelling and detachment. *n* = 7 transwell replicates across three independent dhBMEC differentiations. Statistical significance was calculated by a student's paired t-test. *** $p < 0.001$, ** $p < 0.01$. Data presented as means ± SEM.

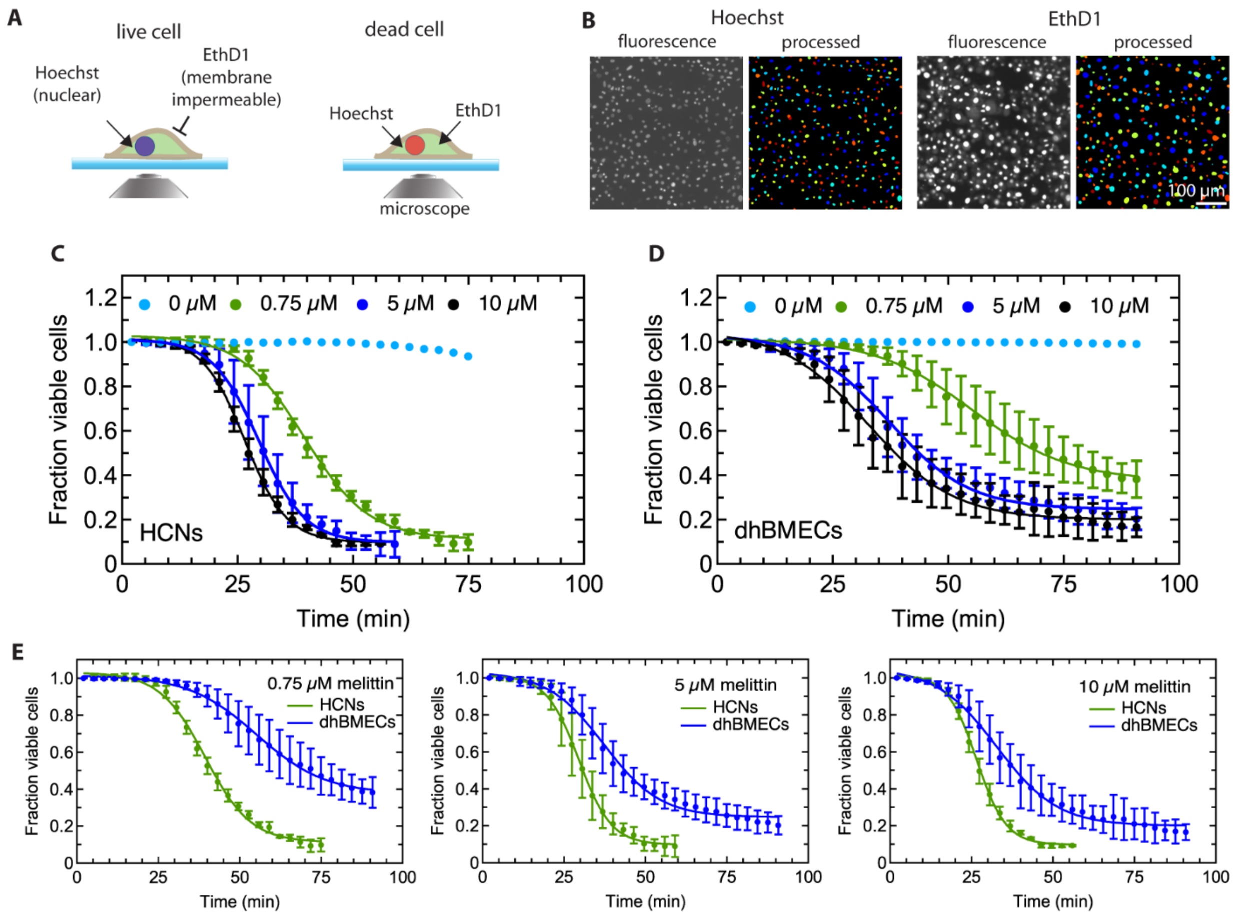


**Fig. S2.** Concentration-dependent melittin-induced cytotoxicity on derived human BMEC-like cells (dhBMECs) and human cortical neuron cells (HCNs). **(A)** A schematic of a cytotoxicity assay. Cells were incubated with Ethidium homodimer-1 (EthD1) dye, which entered cells with permeabilized membranes, and Hoechst dye, which stained nuclei of all cells. The cells were incubated with different melittin concentration together with the assay dyes and imaged using epifluorescence microscope. **(B)** Examples of fluorescence images and processed images to identify the number of EhtD1-positive and Hoechst-positive cells. These images are cropped portions of larger images that were used for the analysis in Cell Profiler [39]. Cell viability was measured by the cell entry of membrane-impermeable Ethidium homodimer-1 dye. **(C)** The effect of exposure of different melittin concentrations on HCN viability. **(D)** The effect of exposure of different melittin concentrations on dhBMEC viability. **(E)** Direct comparison of HCN and dhBMEC viability for different melittin concentrations. Data presented as means ± SEM. $n = 5$ experiments for dhBMECs and $n = 3$ experiments for HCNs.

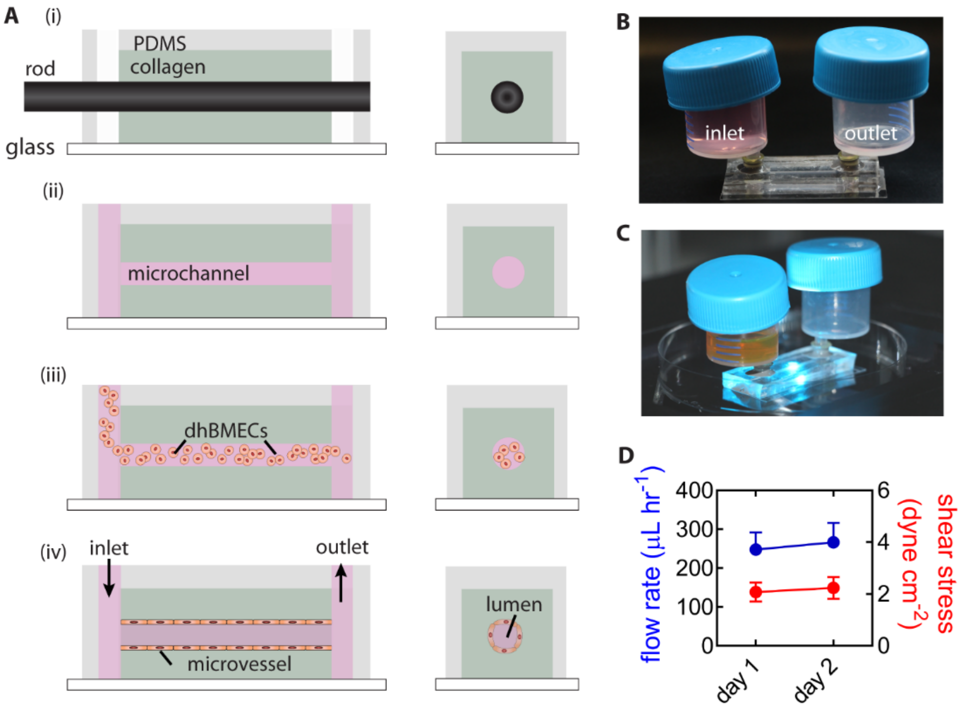


**Fig. S3**. Fabrication and perfusion of tissue-engineered BBB microvessels. **(A)** Fabrication schematic: (i) rods are suspended within 7 mg $mL^{-1}$ type I collagen hydrogels held within rectangular chambers patterned in PDMS, (ii) rod removal leaves behind a 150 μm diameter channel that is stiffened using treatment with 20 mM genipin, (iii) dhBMECs are seeded into channel, and (iv) channels are perfused at approximately 2 dyne $cm^{-2}$. **(B)** Image of flow system comprised on inlet and outlet medium reservoirs. **(C)** Image of device undergoing live-cell imaging on microscope. **(D)** Quantification of flow rates and shear stress over two days after seeding (day 0) (*n* = 14 microvessels). Data presented as means ± SEM.

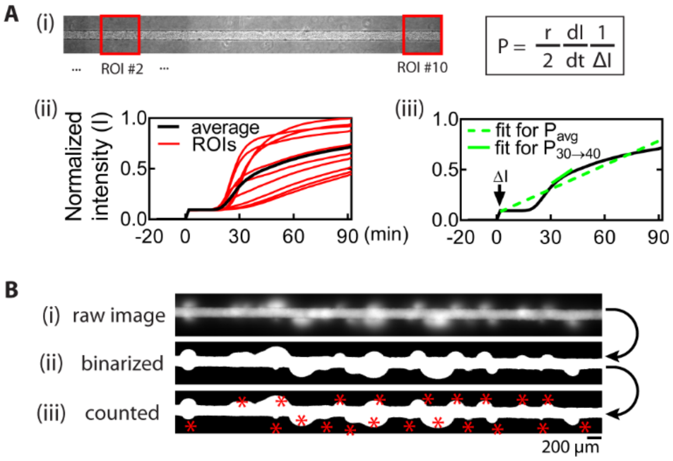


**Fig. S4.** Analysis of BBB opening and recovery within tissue-engineered BBB microvessels. **(A)** 6 mm by 0.6 mm images are sectioned into ten adjacent squares (ROIs #1-10). The normalized fluorescence intensity is plotted over time. Average permeability is calculated using the line of best fit from t = 0 to t = 90 minutes, while instantaneous permeability is calculated using the line of best fit for 10-minute periods for each ROI. **(B)** To quantify focal leaks, raw 500 kDa dextran images were binarized in ImageJ before manual counting of events. Red asterisks represent counted focal leaks.

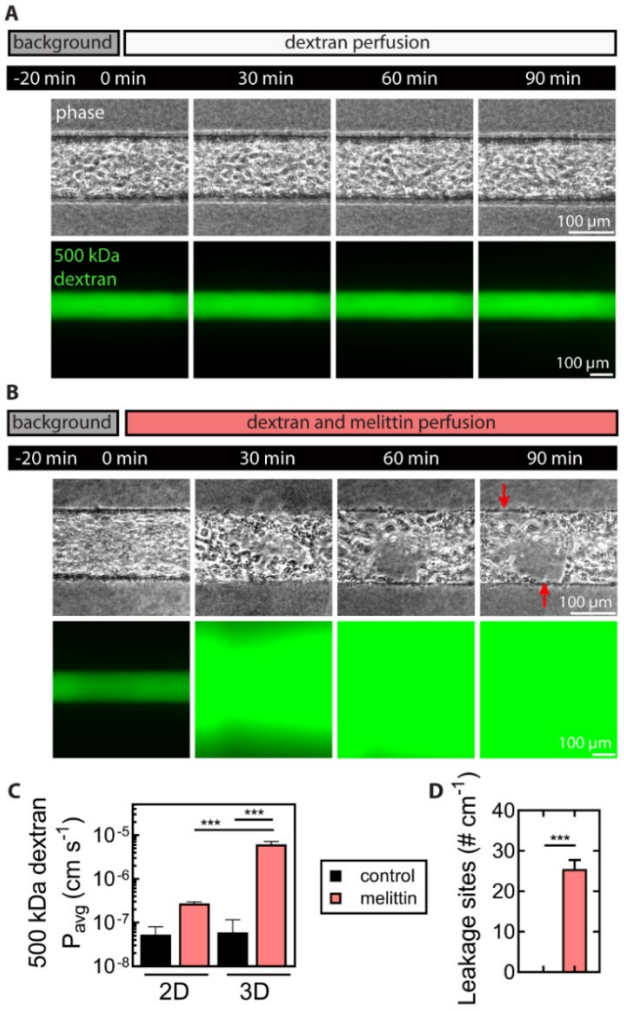


**Fig. S5.** Melittin-induced BBB disruption within tissue-engineered BBB microvessels. Experimental imaging protocol: microvessels were perfused with dextran (A; control) or dextran and 5 µM melittin (B; melittin-exposed) for 90 minutes. **(A)** Control phase contrast and 500 kDa dextran fluorescence time course. A single representative ROI is shown at times 0, 30, 60 and 90 minutes. Microvessel structure remains stable and 500 kDa dextran displays negligible permeability. **(B)** Melittin-exposed phase contrast and 500 kDa dextran fluorescence time course. Formation of defects in the endothelium (red arrows) and high 500 kDa dextran permeability are observed. **(C)** Comparison of 500 kDa dextran permeability in a 2D transwell assay and in BBB microvessels. **(D)** The leakage site density in control and melittin-treated microvessels. $n = 3$ microvessels for each condition. Statistical significance was calculated by one-way ANOVA with Tukey's post hoc test and a student's unpaired t-test. ***$p < 0.001$. Data presented as means ± SEM.

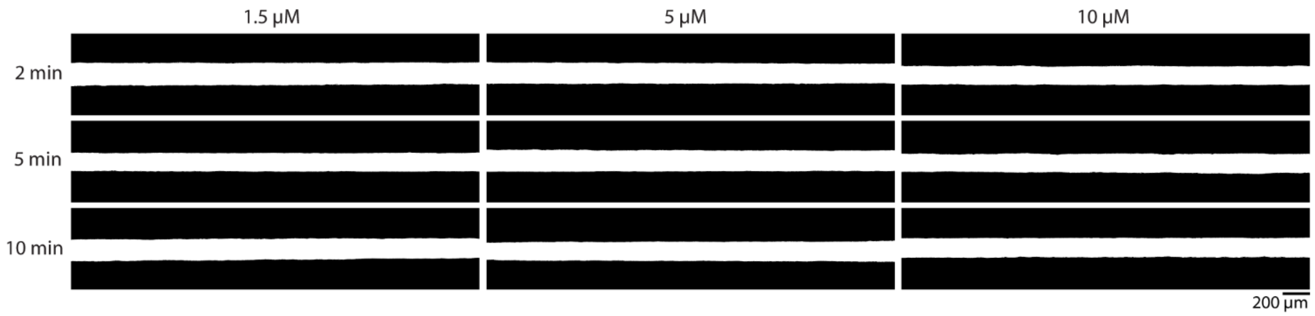


**Fig. S6**. Reversibility of BBB opening 24 hours after melittin exposure. Thresholding of 500 kDa dextran fluorescence images collected 24 hours after melittin exposure. Permeability is reversible 24 hours after the exposure, as 500 kDa dextran remains confined to lumen. Sustained disruption of microvessel barrier would be immediately visible as plumes of dye entering the hydrogel.

Melittin-carboxy
Chemical Formula: $C_{131}H_{228}N_{38}O_{32}$
Exact Mass: 2845.74
Molecular Weight: 2847.50

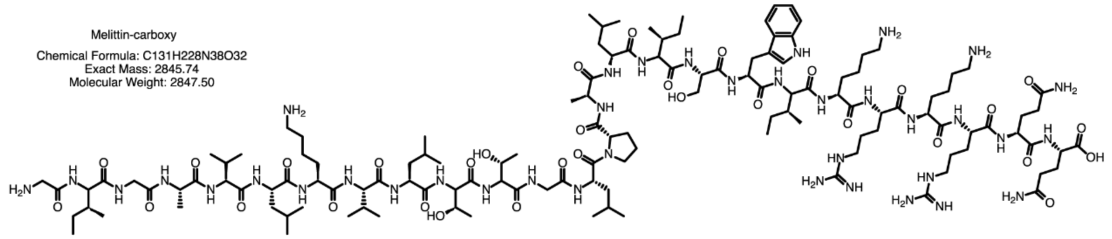

Melittin-caboxyamide
Chemical Formula: $C_{131}H_{229}N_{39}O_{31}$
Exact Mass: 2844.75
Molecular Weight: 2846.52

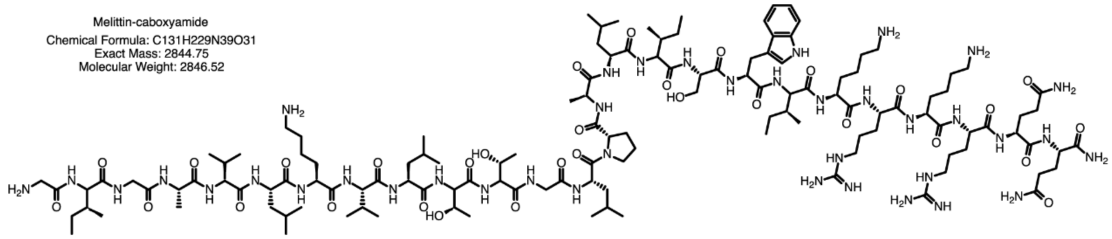

MelP3
Chemical Formula: $C_{127}H_{220}N_{38}O_{30}$
Exact Mass: 2757.69
Molecular Weight: 2759.39

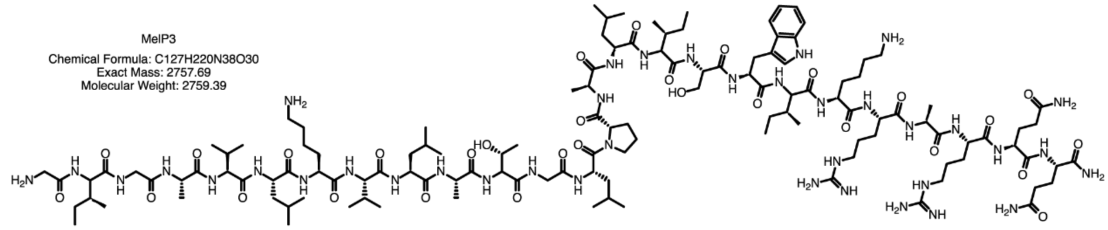

MelP5
Chemical Formula: $C_{124}H_{212}N_{32}O_{30}$
Exact Mass: 2629.60
Molecular Weight: 2631.25

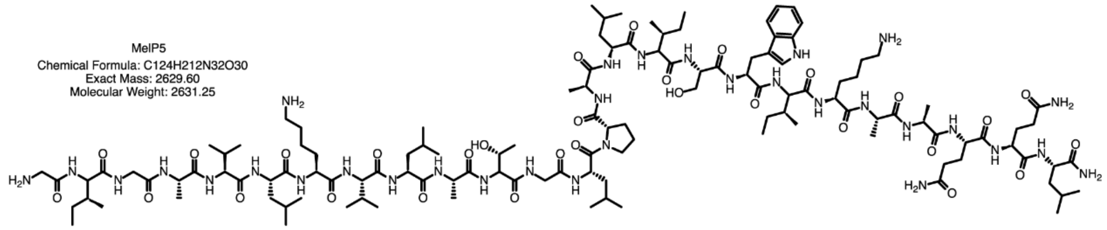

Mel-Scramble
Chemical Formula: $C_{131}H_{229}N_{39}O_{31}$
Exact Mass: 2844.75
Molecular Weight: 2846.52

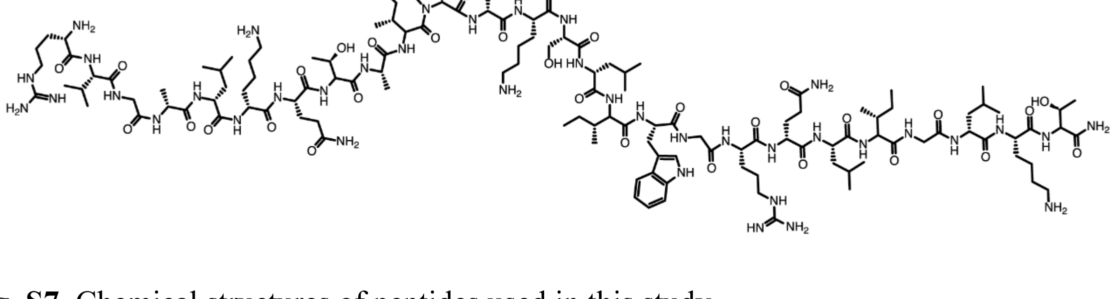

**Fig. S7.** Chemical structures of peptides used in this study.

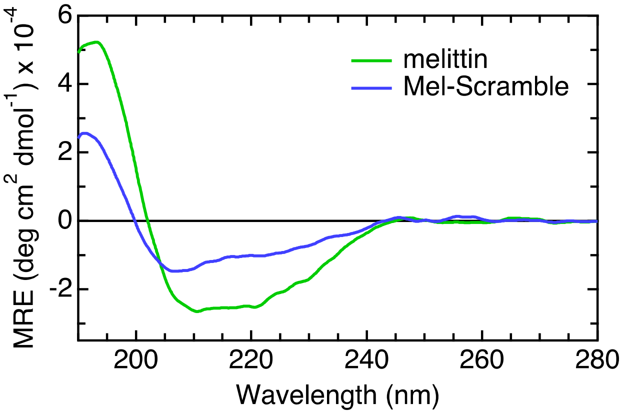


**Fig. S8.** Scrambled melittin (Mel-Scramble) has lower helicity than melittin. Representative circular dichroism spectra of 5 μM melittin and scrambled melittin in 20 mM sodium phosphate buffer with 20 mM SDS, pH 7.4. Spectra were measured 5 times and averaged. Blank buffer spectrum was subtracted, and the spectra were smoothened using a 20-point box algorithm (Igor Pro 7.08; Wavemetrics, Portland, OR, USA).

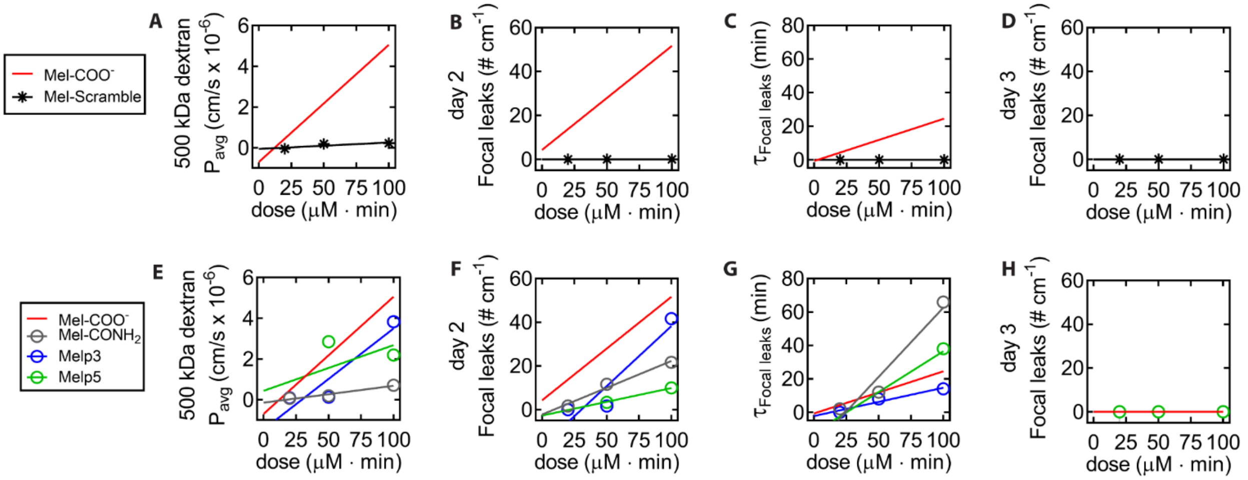


**Fig. S9**. Melittin variants but not scrambled melittin induce reversible BBB opening. **(A-D)** Comparison of BBB opening metrics between scrambled melittin (Mel-Scramble) and melittin (red line) for different doses (25, 50 and 100 μM·min): **(A)** average 500 kDa dextran permeability, **(B)** focal leak density, **(C)** focal leak reversibility constant, and **(D)** focal leak density one day after melittin exposure. **(E-H)** Comparison of BBB opening metrics between melittin variants (symbols) and melittin (red line) for different doses (25, 50 and 100 μM·min): **(E)** average 500 kDa dextran permeability, **(F)** focal leak density, **(G)** focal leak reversibility constant, and **(H)** focal leak density one day after melittin exposure. Total of *n* = 12 microvessels exposed to 25, 50 and 100 μM·min Mel-Scramble, Mel-$CONH_2$, MelP3, and MelP5. Line of best fit for Mel-$COO^-$ data is shown in red as a comparison.

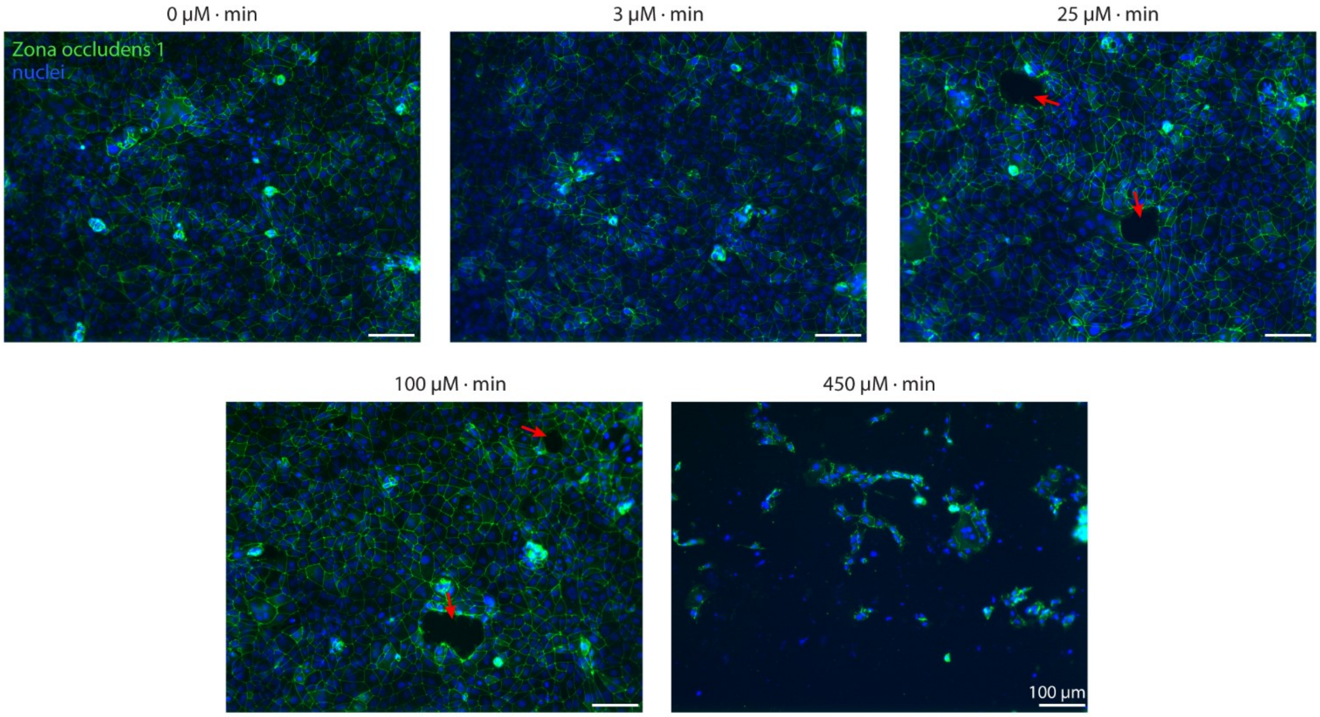


**Fig. S10.** Immunofluorescence imaging of zona occludens-1 (ZO1) junctions after melittin exposure. We do not observe a large effect of melittin on ZO1 expression after 3 μm·min (1.5 μM melittin, 2 min incubation), 25 μM·min (5 μM melittin, 5 min incubation) and 100 μM · min (10 μM melittin, 10 min incubation) melittin treatment. However, melittin treatment is associated with the detachment of cells from the monolayer (indicated by red arrows). Melittin dose of 450 μM·min (5 μM melittin, 90 min incubation) results in a significant cell detachment. Representative images are shown.

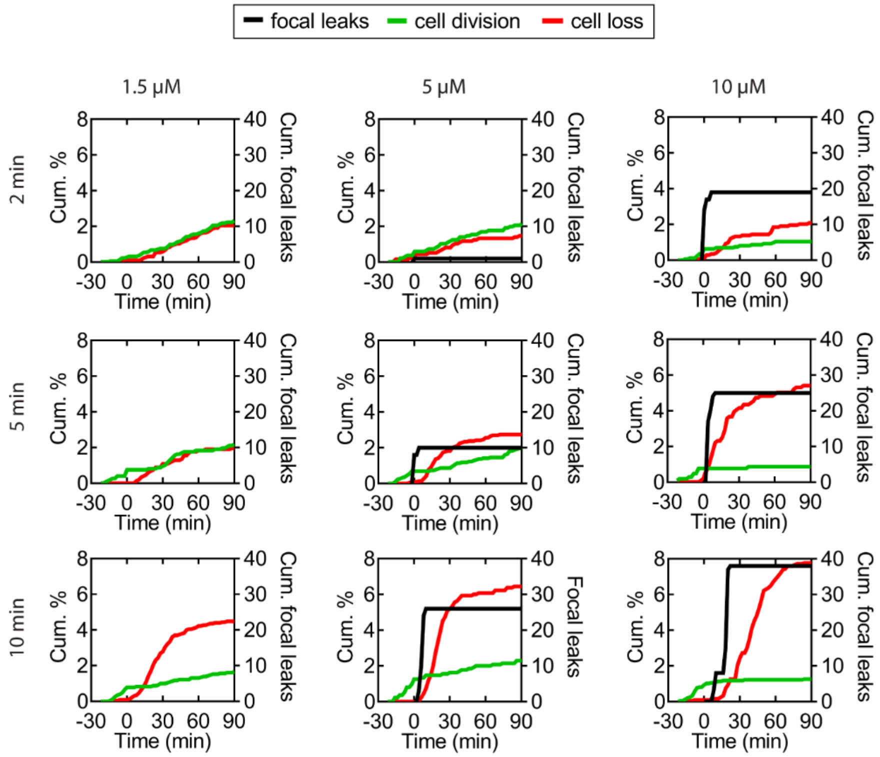


**Fig. S11.** Time course analysis of dhBMEC behavior. Plots of cumulative cell divisions, cell loss, and focal leaks for melittin doses. The left y-axis represents cumulative % of all dhBMECs gained (green line) or lost (red line). The right y-axis represents cumulative number of focal leaks (black line). Results demonstrate that cell loss is observed after focal leak formation. Each graph represents a single melittin dose ($n$ = 9 total).

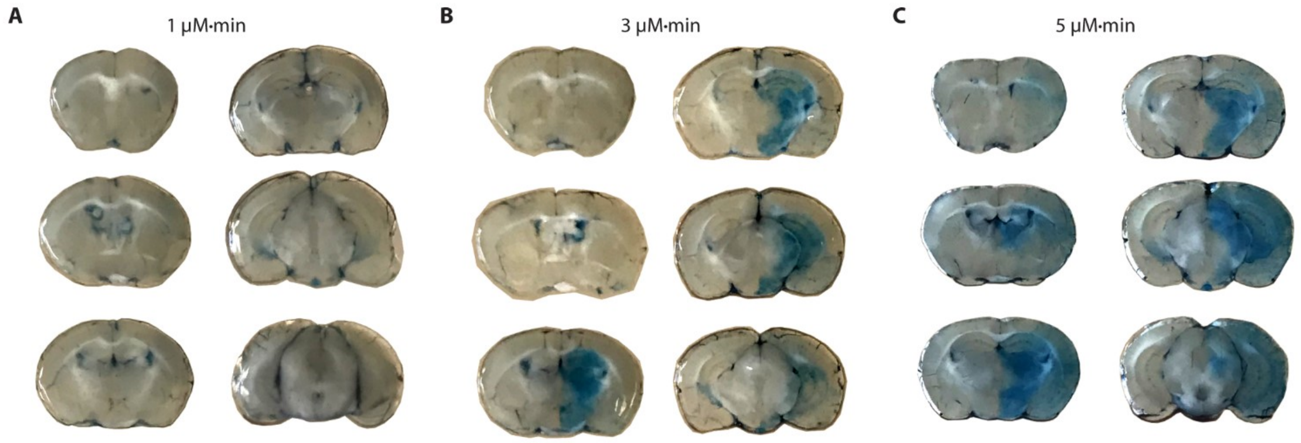


**Fig. S12**. Evans Blue leakage assay to determine minimum effective dose. Representative images are shown of Evans Blue leakage across six mouse brain slices following **(A)** 1 μM·min, **(B)** 3 μM·min, and **(C)** 5 μM·min melittin exposure. For each dose at least $n$ = 3 mice were analyzed.

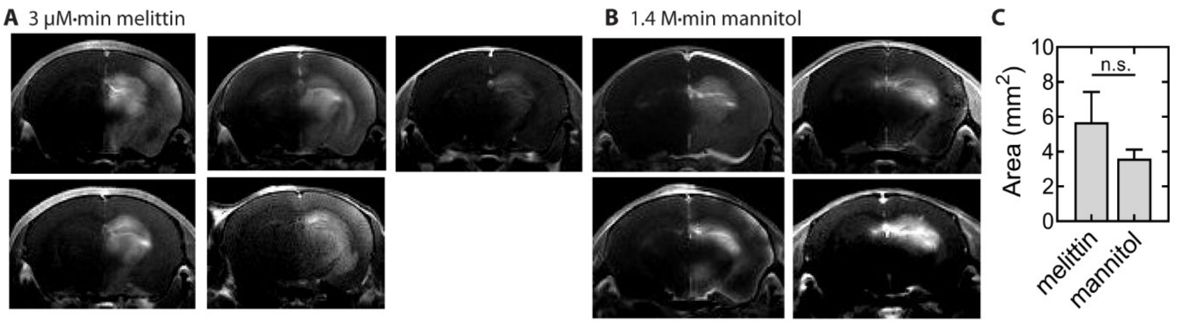


**Fig. S13**. Comparison of melittin- and mannitol-induced BBB opening. **(A-B)** Patterns of gadolinium distribution following 3 μM·min melittin and 1.4 M·min mannitol. Similar variability in BBB opening is observed. Imaging and administration protocols were standardized for both agents; each image represents a unique mouse. **(C)** Comparison of BBB opening area between approaches. *n* = 5 for melittin and *n* = 4 for mannitol. Statistical significance was calculated by a student's unpaired t-test. Data presented as means ± SEM.

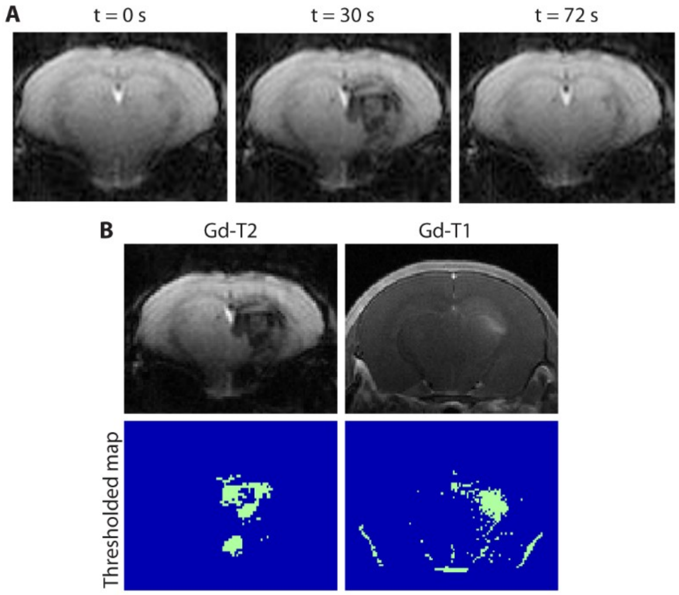


**Fig. S14**. Perfusion territory of intraarterial injections. **(A)** Dynamic Gd-T2 MRI images of mouse brain before the intraarterial infusion of Gd, and 35 s and 75 s after the infusion at a rate of 150 μL $min^{-1}$ showed that mainly deep brain structures (hippocampus particularly) are perfused. **(B)** Thresholding of Gd-T2 pre-melittin and Gd-T1 post-melittin to correlate perfusion and BBB opening territories.

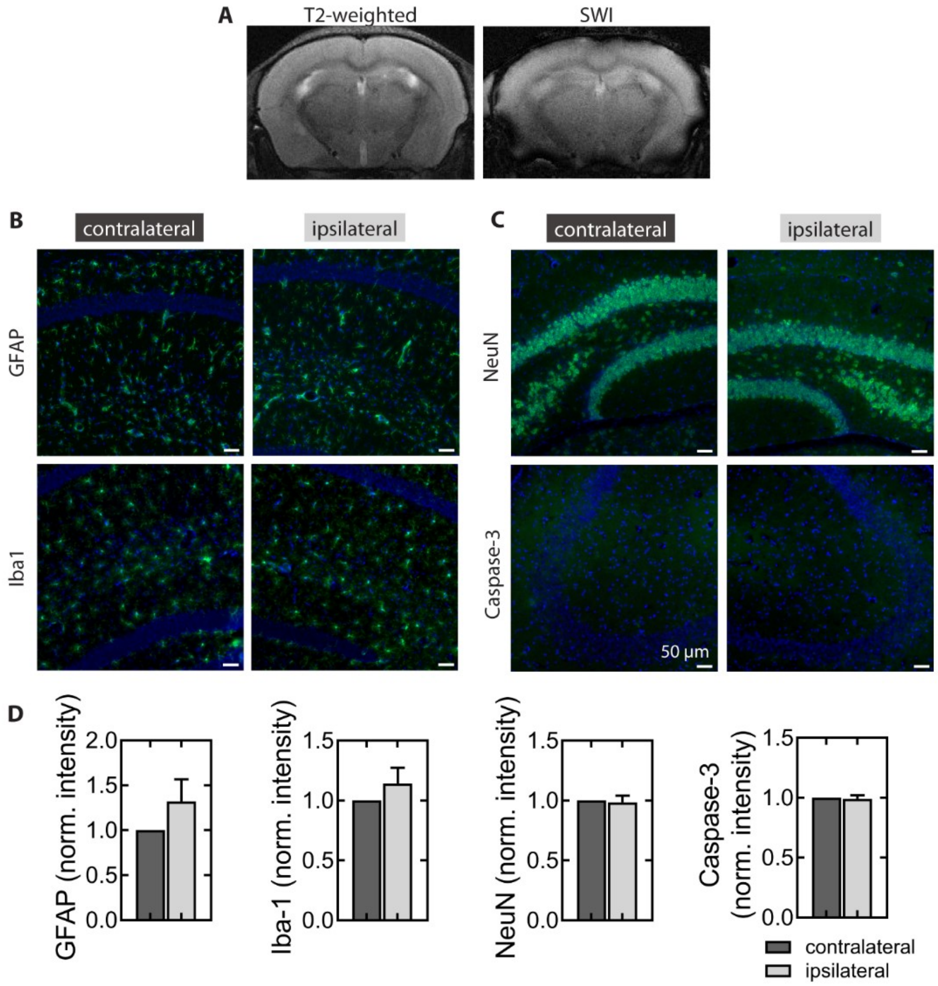


**Fig. S15**. MRI and histological assessment 24 hours following melittin-induced BBB opening. **(A)** T2-weighted and SWI images 24 hours after BBB opening (BBBO) showed no sign of brain damage or microhemorrhage, respectively. **(B)** Immunohistochemical detection of neuroinflammation markers glial fibrillary acidic protein (GFAP) and ionized calcium binding adaptor molecule 1 (Iba1) in the contralateral and ipsilateral BBBO region. **(C)** Immunohistochemical detection of damage markers using a neuronal nuclear antigen (NeuN) and apoptotic marker (caspase-3) in the contralateral and ipsilateral BBBO region. **(D)** Comparison of normalized intensity of immunohistochemical images between the contralateral (control) and ipsilateral BBBO region. Representative images are shown. $n = 3$ mice for each imaging technique or stain. Statistical significance was calculated by a student's paired t-test. Data presented as means ± SEM.

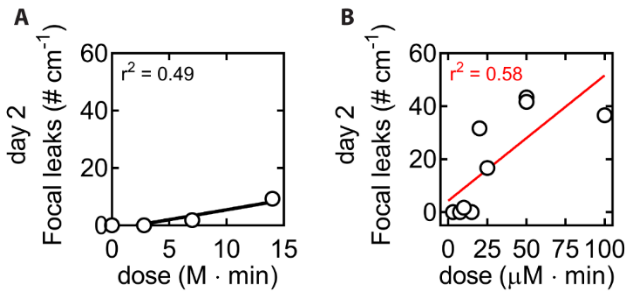


**Fig. S16.** Retrospective comparison of mannitol and melittin-induced BBBO within tissue-engineered microvessels. **(A)** Focal leak density over mannitol dose. **(B)** Focal leak density over melittin dose.

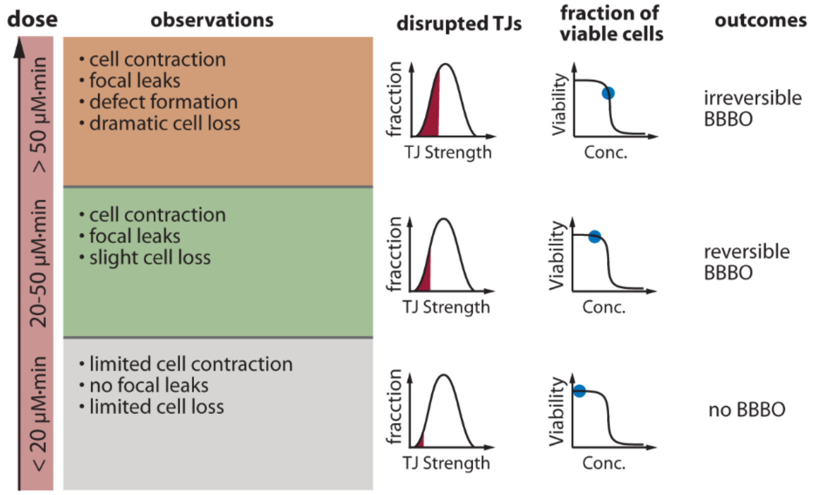


**Fig. S17.** Mechanisms of melittin-induced BBBO. We summarize observations and outcomes of various melittin doses on BBB microvessels. At low doses (below 20 μM·min in microvessels), endothelial cells are able to dynamically retain cell-cell contacts during cell division and cell loss thereby maintaining barrier function and preventing focal leaks. This process is not associated with disruption of cell-cell junctions or loss of viability. At intermediate doses (20 μM·min – 50 μM·min in microvessels), the tensile forces resulting from cell contraction are sufficiently large to induce local disruption of weak cell junctions resulting in the formation of focal leaks. The density of contraction and disruption events is sufficiently low that they are isolated, and recovery of normal barrier function occurs rapidly. At high doses (> 50 μM·min in microvessels) the density of contraction events is sufficiently high that clusters of disrupted cell-cell junctions results in dramatic cell loss.

**Table S1.** Amino acid sequences of melittin variants.

| Peptide | Sequence | MW (Da) |
|---|---|---|
| Melittin-$COO^-$ | GIGAVLKVLTTGLPALISWIKRKRQQ-OH | 2847.5 |
| Melittin-$CONH_2$ [a] | GIGAVLKVLTTGLPALISWIKRKRQQ-$NH_2$ | 2846.5 |
| MelP3 [a,b] | GIGAVLKVL**A**TGLPALISWIKR**A**RQQ-$NH_2$ | 2759.4 |
| MelP5 [a,b] | GIGAVLKVL**A**TGLPALISWIK**AAQ**Q**L**-$NH_2$ | 2631.3 |
| Mel-Scramble [a,c] | RVGALKQTAIPVKSLIWGRQLIGLKT- $NH_2$ | 2846.5 |

[a] Variants contain carboxyamide ($CONH_2$) C-terminus.

[b] Variants discovered in a high-throughput screen for the ability to have high pore-forming activity in lipid bilayers [26]. Amino acids different from wild-type melittin sequence are bolded.

[c] Scrambled melittin variant designed to have low primary and secondary amphipathicity.

**Table S2.** Primary antibodies used in this study.

| Antigen | Description | Vendor | Cat. No. | Dilution |
|---|---|---|---|---|
| Zona occludens-1 | rabbit polyclonal | Life Technologies | 40-2200 | 1:200 |
| Claudin-5 | mouse monoclonal | Life Technologies | 35-2500 | 1:100 |
| Occludin | rabbit polyclonal | Life Technologies | 40-4700 | 1:100 |
| GFAP | rabbit polyclonal | Dako | Z0334 | 1:250 |
| Iba1 | rabbit polyclonal | Wako | 019-19741 | 1:250 |
| Caspase 3 | rabbit polyclonal | Cell Signaling Technology | D175 | 1:400 |
| NeuN | rabbit monoclonal | Cell Signaling Technology | D3S3I | 1:100 |

**Movies:**

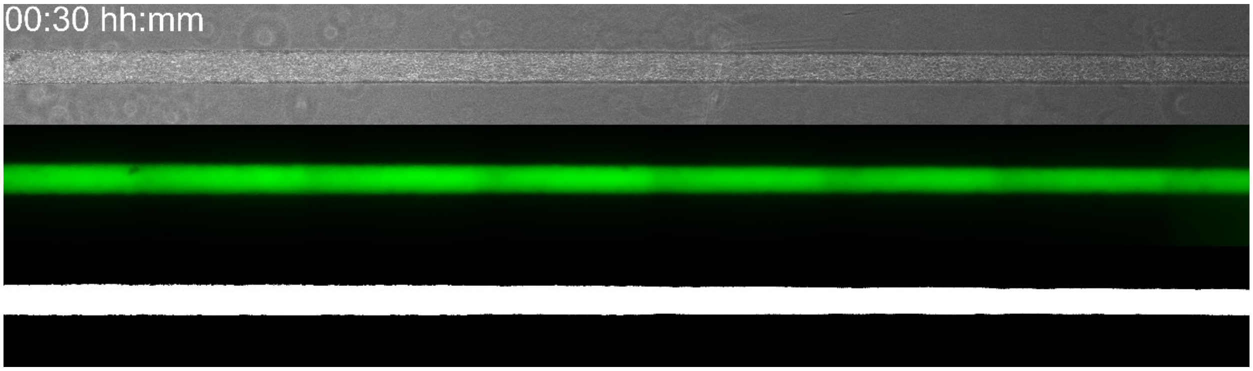


**Movie S1**. BBB microvessels display negligible permeability to 500 kDa dextran. Timelapse images over 90 minutes of (top) microvessel midplane, (middle) 500 kDa dextran fluorescence, (bottom) thresholding of 500 kDa dextran fluorescence. Microvessel is approximately 150 μm in diameter.

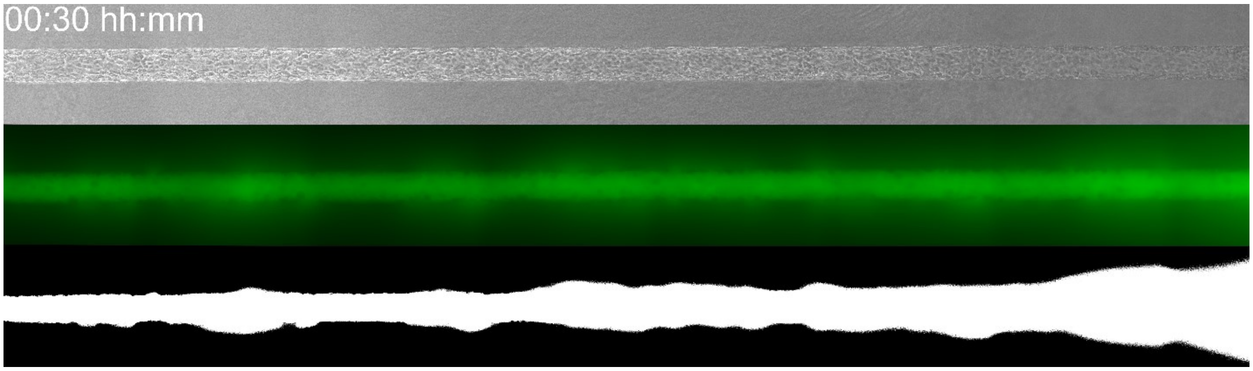


**Movie S2.** BBB disruption during continual perfusion of 5 μM melittin. Timelapse images over 90 minutes of (top) microvessel midplane, (middle) 500 kDa dextran fluorescence, (bottom) thresholding of 500 kDa dextran fluorescence. Focus on the midplane shifts during imaging due to loss of BMECs causing microscopy auto-focus to fail. Microvessel is approximately 150 μm in diameter.

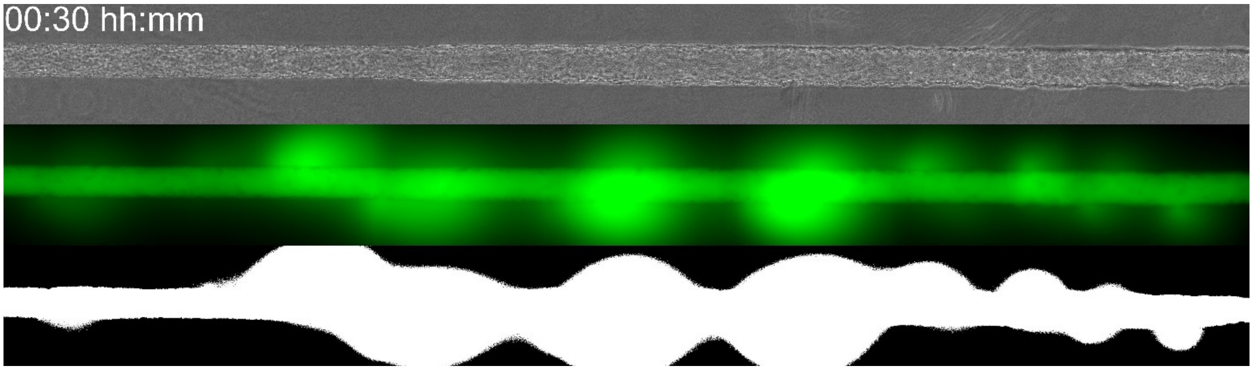


**Movie S3.** Reversible BBB opening with a 10 min 5 μM melittin dose. Timelapse images over 90 minutes of (top) microvessel midplane, (middle) 500 kDa dextran fluorescence, (bottom) thresholding of 500 kDa dextran fluorescence. Transient BBB opening is observed as focal leaks of 500 kDa dextran entering the ECM (from t = 4 to 34 minutes) followed by focal leak cessation. Although macroscopic structure of microvessel is not altered, increased turnover of cells is observed as balled up cells flowing through the microvessel lumen. Microvessel is approximately 150 μm in diameter.

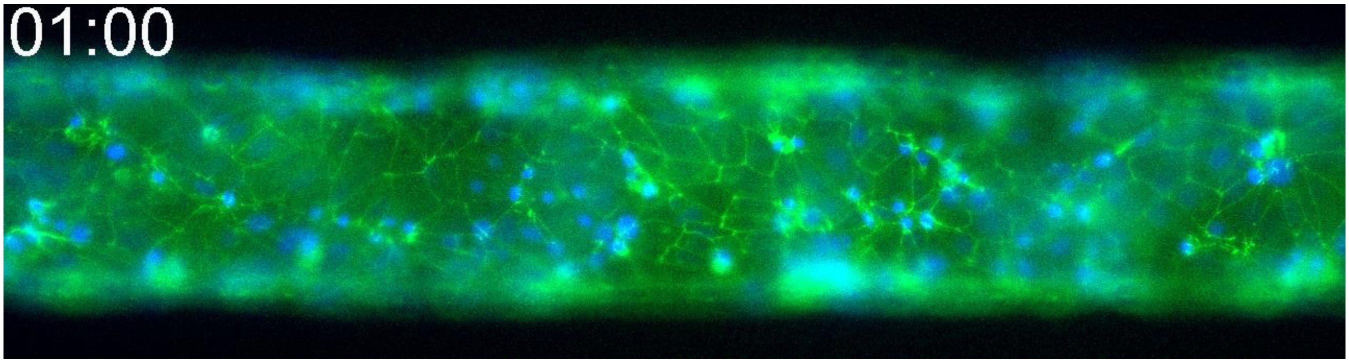


**Movie S4.** Imaging transient BBB opening using microvessels comprised of zona-occludens-1 fluorescently-tagged dhBMECs. Exposure to melittin induces cell contraction and condensation of zona occludens-1 to an intracellular localization. Green = zona occludens-1, blue = DAPI (nuclei). Microvessel is approximately 150 μm in diameter.

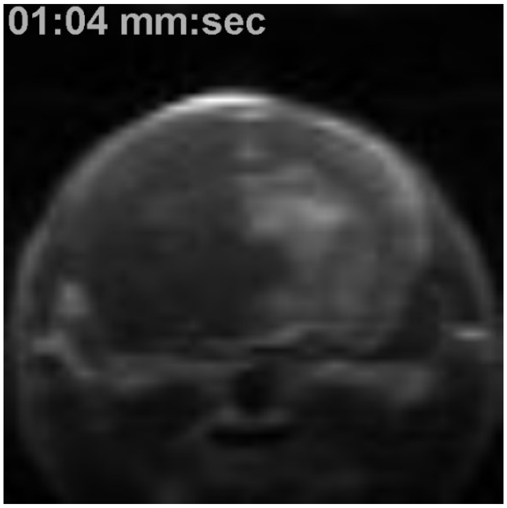


**Movie S5.** Real-time imaging of BBB opening of mouse brain using 3 μM melittin for 1 minute under gadolinium enhancement on T1-weighted imaging. Shown is imaging of the hippocampus. Injection began at t = 0 seconds (first image frame).
